# Ancient intraflagellar transport machinery controls unique spatial distribution of phototropin in an evolutionary important non-flagellated vegetative stage of terrestrial alga

**DOI:** 10.64898/2026.02.05.698958

**Authors:** Manisha, Rajani Singh, Sunita Sharma, Suneel Kateriya

**Affiliations:** Laboratory of Optobiotechnology, School of Biotechnology, Jawaharlal Nehru University, New Delhi 110067, India; Department of Cellular Biology, University of Georgia, Athens 30602, Georgia, USA

## Abstract

Intraflagellar transport (IFT) is a conserved trafficking system in eukaryotes that moves proteins along microtubules. It is best known for its essential role in building and maintaining cilia and flagella. Intriguingly, several IFT components are still found in organisms that no longer possess flagella, raising important questions about their original functions and how they may have been repurposed during evolution. The filamentous alga *Klebsormidium nitens*, positioned at the base of the streptophyte lineage, offers a valuable model for exploring this transition. Here, we investigate the IFT machinery in *K. nitens* and its relationship with the blue-light photoreceptor phototropin. Comparative genomic analyses show that key IFT-A and IFT-B components are retained, despite the complete loss of flagella in vegetative state. Cellular detection and immunofluorescence studies revealed the presence and localisation of IFT components, interestingly, their co-localization with phototropin. Notably, IFT-139 and IFT-20 strongly co-localize with phototropin at plasma membrane-associated regions. Phototropin overlapping localization (plasma membrane associated) with conserved phospho-adaptor protein 14-3-3, pointing to a phosphorylation-dependent signaling network. Unlike in *Chlamydomonas reinhardtii*, where these proteins localize to flagella, their interaction in *K. nitens* occurs independently of cilia presence. Together, these results evidenced that IFT components were retained and repurposed early in streptophyte evolution and might support phototropin localization and signalling, revealing an ancestral, non-ciliary role for the IFT system.

## Introduction

The transition of plants from an aquatic to a terrestrial environment represents one of the most transformative events in the evolution of life on Earth. This transition required profound cellular and molecular innovations, particularly in light perception, signal transduction, and cytoskeletal organization, enabling early photosynthetic organisms to cope with fluctuating light conditions, gravity, and desiccation stress (Delwiche & Cooper, 2015; Ishizaki, 2017; Kapoor et al., 2023; López-Pozo et al., 2023). Particularly, light perception is essential for photosynthetic organisms as it plays a central role in regulating growth and development in plants and algae according to changing environmental conditions. This perception is mediated by light-sensitive proteins, photoreceptors, which enable organisms to detect light corresponding to different wavelengths. Among the blue-light photoreceptors, phototropin is a major player in light-induced directional response and other key light-driven adaptive responses such as stomatal opening, chloroplast movement, phototaxis and phototactic responses, leaf expansion and flattening (Christie, 2007; Takemiya et al., 2005). Phototropins are flavoproteins consisting of 2 LOV (Light, oxygen and Voltage) domains at the N-terminus and a kinase domain at the C-terminus, which is responsible for initiating signalling after blue light-induced conformational changes (Hart & Gardner, 2021). They are conserved across most of the land plant lineages and in green algae, suggesting that phototropin-mediated light signalling evolved prior to land colonisation. Charophyte green algae, which are the closest living relatives of land plants, provide a unique evolutionary window to investigate the origins of these innovations. Among them, *Klebsormidium nitens*, a filamentous streptophyte alga, has emerged as a key model system for studying the molecular foundations underlying plant terrestrialization and light adaptive responses (Hori et al., 2014). In the model chlorophyte alga *Chlamydomonas reinhardtii*, phototropin is known to perform various light dependent functions. These include regulation of progression of sexual life cycle, modulation of starch metabolism in response to blue light, regulates expression of genes involved in chlorophyll and carotenoid biosynthetic pathways, in phototaxis and phototactic process and regulation of chemotaxis towards ammonium ion (Im et al., 2006; Yuan et al., 2025). It requires precise subcellular localisation to perform above aforementioned functions. In algae, it is known that intraflagellar transport (IFT) machinery is needed for localisation of phototropin at different organelles such as flagella and eyespot (Huang et al., 2004). As a result, in motile algae, IFT serves as a direct mechanistic bridge between photoreceptor signaling and the intracellular trafficking of proteins. IFT machinery is bidirectional transport system known for trafficking the phototropin along the length of flagella. It is composed of 2 subunits-IFT-A and IFT-B along with motor proteins kinesin and dynein to drive anterograde and retrograde movement respectively (Taschner & Lorentzen, 2016). Although, IFT has been studied primarily in flagellated organisms, growing genomic evidence indicates that many IFT components are also retained in primitive non-flagellated lineages (Hodges et al., 2012; Hori et al., 2014). This observation supports the idea that IFT may be an evolutionarily ancient system involved in intracellular trafficking and scaffolding, existing well before the loss of flagella. In higher plants, phototropins are mainly localized at the plasma membrane, where they sense blue light and trigger downstream signalling pathways. Following light activation, phototropins can dynamically relocalize, showing partial redistribution to the cytosol or to distinct membrane subdomains (Kong et al., 2013; Liscum, 2016). This spatial regulation is controlled through phosphorylation, interactions with regulatory proteins such as 14-3-3 proteins, and vesicle-mediated intracellular trafficking (Reuter et al., 2021). Precise control of phototropin transport and localization is crucial for its roles in key physiological processes, including phototropism, stomatal opening, and chloroplast movement. The presence of phototropin-mediated blue-light signaling in *K. nitens* (Manisha et al., 2025), therefore, raises an important evolutionary question: how is phototropin spatially organized in the absence of flagella, does its localization resemble that observed in higher plants, and have components of the IFT machinery been retained and repurposed to support photoreceptor localization and signalling? In this study, we identified the photoreceptor machinery in *K. nitens* mainly focussing on the blue light sensing protein, phototropin. Importantly, we report the first detailed characterization of IFT machinery in terrestrial alga, *K. nitens* in vegetative state and examine its spatial relationship with the blue-light photoreceptor phototropin. By combining bioinformatic analyses, biochemical, molecular and cellular approaches, we investigated the subcellular localization of key IFT components and assess their colocalization with phototropin.

## Materials and methods

Bioinformatic analysis-For homology analysis, protein sequence of different photoreceptors present in *Chlamydomonas reinhardtii* and *Arabidopsis thaliana* were used as queries and blasted against *K. nitens* genome database using NCBI and Phytozome. Further, for identification of IFT components, *C. reinhardtii* and *Homo sapiens* protein sequence were used as reference for BLAST searches in *K. nitens* genome https://phycocosm.jgi.doe.gov/Klenit1_1/Klenit1_1.home.html present in Phycocosm database and NCBI. Using domain prediction tools CD search https://www.ncbi.nlm.nih.gov/Structure/cdd/wrpsb.cgi and CDART https://www.ncbi.nlm.nih.gov/Structure/lexington/lexington.cgi candidate proteins were further analysed for conserved domains. Protein sequences retrieved from NCBI and Phycocosm were submitted to STRING database (Szklarczyk et al., 2025) for protein protein network predictions, and later it was visualised and analysed using Cytoscape V 3.10.3 (Shannon et al., 2003).

### Cell culture and growth conditions

*Klebsormidium nitens* cells were sourced from UTEX Culture Collection of Algae, the University of Texas, Austin (https://utex.org/). *Klebsormidium nitens* was grown and cultured in Pentose Peptone media (PPM). The PPM media constitute different components-NaNO_3_, CaCl_2_.2H_2_O, MgSO_4_.7H_2_O, K_2_HPO_4_, KH_2_PO_4_, NaCl and Protease peptone. Cultures were grown under 14 h light and 10 h dark photoperiod at a constant temperature of 22 °C with continuous shaking at 120 rpm. Cells were maintained in solid PPM containing 1.5% agar and grown in liquid PPM medium under white LED light with intensity of approx. 2500 LUX.

### Cells harvesting, protein extraction and immunoblotting

Cells were grown until reaches the optical density (OD) 0.3-0.4 and then harvested using centrifugation at 10,000 rpm. The harvested cells were resuspended in lysis buffer containing 1X Phosphate Buffer Saline (PBS), phenylmethylsulphonyl fluoride (200 µM PMSF) and 0.1% protease inhibitor cocktail (PIC). Further, process of lysis was done using sonication method with 7 sec ON/OFF pulse for 15 cycles at 30% amplitude. At this point, TCL is collected. Total cell lysate was further centrifuged at 13000 rpm for 30 min. Then the obtained supernatant along with TCL fraction was mixed with 2X laemmli buffer and denaturated at 65°C for 30 min. For western blot analysis, 100 µg of protein was used, protein concentration was estimated using Bradford reagent. Equal amount of proteins were loaded in the each well and resolved on 8-15% SDS-PAGE. It is then transferred to nitrocellulose membrane. Membrane was blocked overnight using 5% skimmed milk with PBST (PBS+0.1% tween-20) at 4°C. Next day, the blocked membrane was incubated with lab raised antigen specific primary antibody for 1 h at room temperature, followed by three washes with 1X PBST for 10 min. Then membrane was incubated with HRP conjugated secondary antibody for 1 h at room temperature on rotor shaker, followed by thrice washing of membrane using 1XPBST for 15 min. Membrane was developed using enhanced chemiluminescence reagent Super signal West Pico Plus (Thermo Scientific) (Awasthi et al., 2016).

### Immunofluorescence microscopy

For cellular localization, *Klebsormidium* cells grown till O.D-0.3-0.4 were centrifuged at the end of the dark phase at 13000 rpm for 30 min. The cell pellet was resuspended in 1XPBS and used for seeding. Coverslips were coated with 0.01% (w/v) poly-L-lysine and incubated at room temperature for 30 min. Cells were seeded on the coverslips treated with poly-L-lysine and incubated for 30 min at room temperature. Fixing of the cells was done using 4% freshly prepared formaldehyde solution in 1XPBS for 20 min. To permeabilise the cells, coverslips were dipped into ice cold absolute ethanol at -20°C for 20 min. Ethanol was removed and cells were incubated with 250 mM NaCl prepared in 1XPBS for 15 min at room temperature. Following rehydration with NaCl, coverslips were rinsed with 1XPBST (1XPBS+0.5% Triton X) four times. Then the cells were incubated with diluted primary antibodies (dilution 1:250-1000) for overnight at 4°C. Following overnight incubation, cells were washed with 1XPBST four times with 1 min each wash. After that coverslips were incubated with fluorophore conjugated secondary antibodies (dilution 1:1000) for 1h at 37°C, followed by washing with 1XPBST for 4 times in 5 min. Further three washes were done using 1XPBS to remove the nonspecific binding of antibodies (Awasthi et al., 2016). Before imaging, coverslips were mounted to glass slides using SlowFadeTM Gold antifade reagent. The mounted coverslip was examine using microscope, Leica STELLARIS (STED).

### RNA isolation, cDNA synthesis and qRT-PCR

To compare the expression levels of photoreceptor and IFT components under different light conditions, real-time quantitative PCR was performed. *K*.*nitens* cultures were exposed to different light conditions (∼ 2500 LUX) for 2 h with continuous shaking at 120 rpm, followed by harvesting through centrifugation at 13000 rpm. Total RNA was extracted from different samples using Trizol method. Following RNA extraction, cDNA synthesis was done using verso DNA synthesis kit (Thermoscientific) in accordance to manufacturer protocol. The qRT-PCR was done following method adapted from (Manisha et al., 2025). Tubulin was used as internal reference gene for normalization. The dissociation curve analysis was also performed to ensure the accuracy of the experiment. The list of the primers used in this study is given in the table S1. Delta-delta Ct method was used to calculate the change in relative expression pattern (Livak & Schmittgen, 2001).

## Results and Discussion

### Phototropin operates within a signalling network in *Klebsormidium nitens* is evolutionarily conserved, yet little rewired

Phototropin showed extensive connectivity with a wide range of signaling and regulatory proteins, underscoring its role as a central integrator of light-responsive pathways. Notably, phototropin showed interaction with multiple photoreceptors, including UVR8, rhodopsin-like proteins, and members of the photolyase/cryptochrome family (Sharma et al., 2021), suggesting coordinated cross-talk between blue- and UV-light perception systems (Fig. 1). The interaction network also revealed strong associations between phototropin and key signaling regulators such as 14-3-3 proteins, AGC family kinases, serine/threonine protein phosphatases, and Ras-related small GTPases. These interactions highlight phosphorylation-dependent regulation and GTPase-mediated signalling as important downstream components of phototropin function. Furthermore, the presence of REM2- and Rab-like small GTPases within the phototropin interactome points to a potential connection between light perception, membrane trafficking, and cytoskeletal dynamics (Sharma et al., 2021); Sharma et al., 2025).

**Fig. 1:**
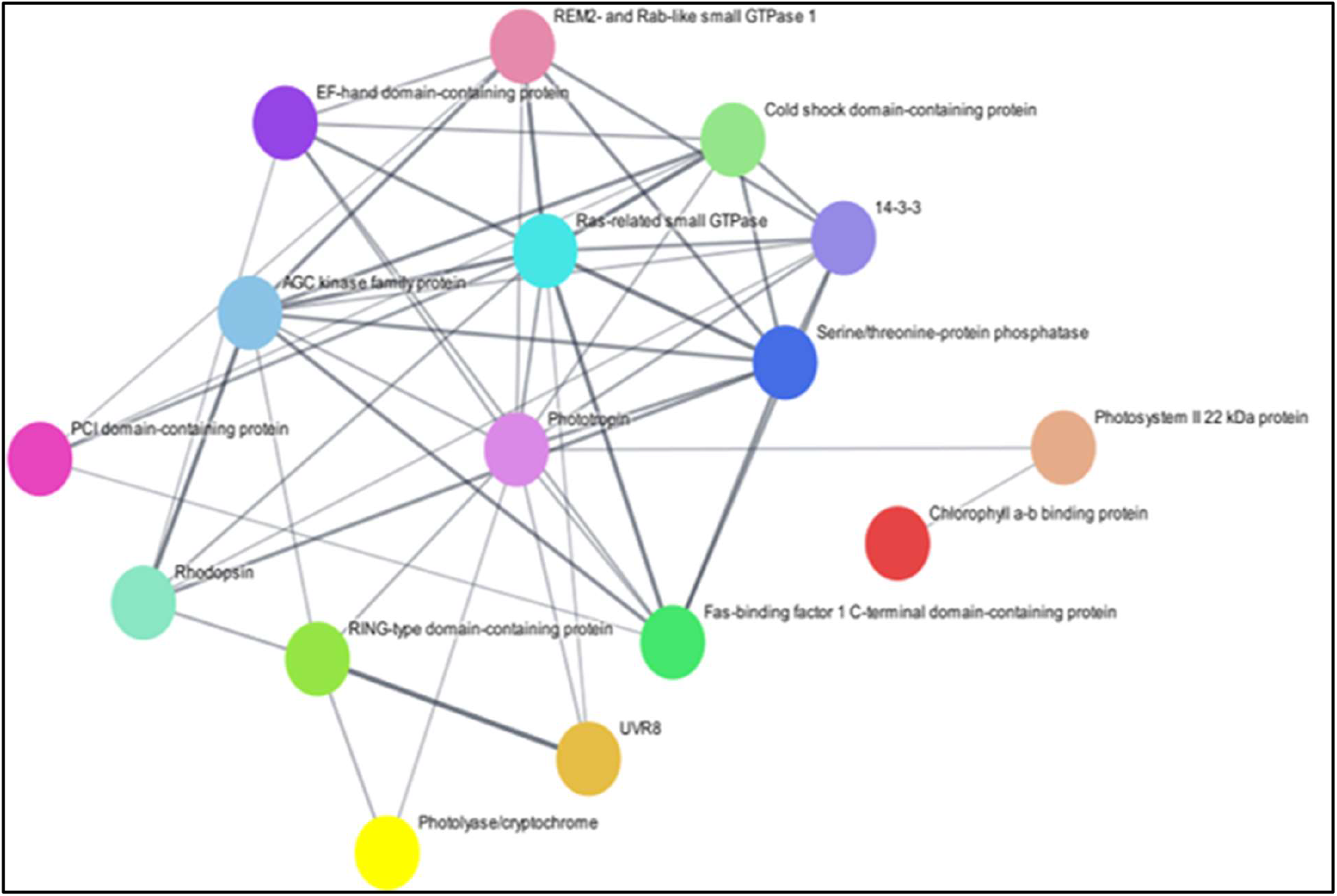
Predicted protein-protein interaction network and cellular detection of phototropin and its regulatory protein 14-3-3 of *Klebsormidium nitens*. Interactome prediction showing the interaction of phototropin (central node) with various regulatory and signalling proteins such as 14-3-3 protein, different phosphates and kinase, and other photoreceptors associated with light signaling pathway.

Interestingly, phototropin exhibited a direct interaction with a homolog of cilia and flagella-associated proteins (CFAPs) in *K. nitens* (Fig. 1a). CFAPs, also known as flagellar-associated proteins (FAPs), are well characterized in flagellated organisms, where they play crucial roles in cilia and flagella assembly (Wang et al., 2022). In *C. reinhardtii*, CFAPs contribute to axonemal organization and motility by stabilizing the outer doublet microtubules of the axoneme and by functioning as scaffold proteins that coordinate both structural and regulatory components of the flagellum (Hou et al., 2021; Yanagisawa et al., 2014). Among these proteins, BUG22 represents a conserved CFAP that has been identified not only in flagellated algae, but also in higher plants such as *A. thaliana*, which completely lack motile cilia and centrioles. The persistence of BUG22 homologs in land plants, despite the loss of centrioles and cilia suggests that BUG22 proteins have adopted non-ciliary functions during evolution. Although plants no longer possess flagella, they retain functional microtubule-organizing centers (MTOCs) and highly ordered cortical microtubule arrays. Thereby, BUG22 is proposed to play a role in facilitating interactions between microtubules and the plasma membrane, thereby contributing to the spatial organization and stability of cytoskeletal networks (Laligné et al., 2010). Thus, following the speculation, we infer that CFAP in *K. nitens* might be likely associated with microtubules arrays, thereby playing role in cytoskeleton organization. In the flagellated green alga *C. reinhardtii*, phototropin shows a highly specific subcellular localization to axoneme through IFT (Huang et al., 2004). The interaction of phototropin with axonemal microtubules allows blue-light perception to be spatially coupled with flagellar motility, enabling rapid and precise modulation of flagellar beating during phototactic responses. This axoneme-centered organization of phototropin serves as an important evolutionary reference for understanding how phototropin signalling has been restructured in non -flagellated algae, *K. nitens*.

### Comparative analysis of the domain architecture of intraflagellar transport (IFT) components revealed a strong degree of structural conservation between non-flagellated vegetative state of *Klebsormidium nitens* and the flagellated green alga *Chlamydomonas reinhardtii*

Based on the observations in Fig. 1, we performed the detailed characterization of IFT components in the algae *K. nitens*. At first, using different bioinformatics tools, we identified the different domains of IFT components present in *K. nitens* (Fig. 2a). We found that the core subunits of both the IFT-A and IFT-B complexes in *K. nitens* retain the hallmark domains characteristic of canonical IFT proteins (Fig. 2a). These include multiple WD40 repeats and TPR motifs in IFT-A subunits, as well as coiled-coil (CC), calponin homology (CH), TPR, and GTPase-associated domains within IFT-B components, indicating a conserved molecular framework despite the absence of flagella. In order to discover the role of these components in *K. nitens*, we further compared the domain architecture of IFT components of *K. nitens* with *C. reinhardtii* (Fig. S1, Table S2).

**Fig. 2:**
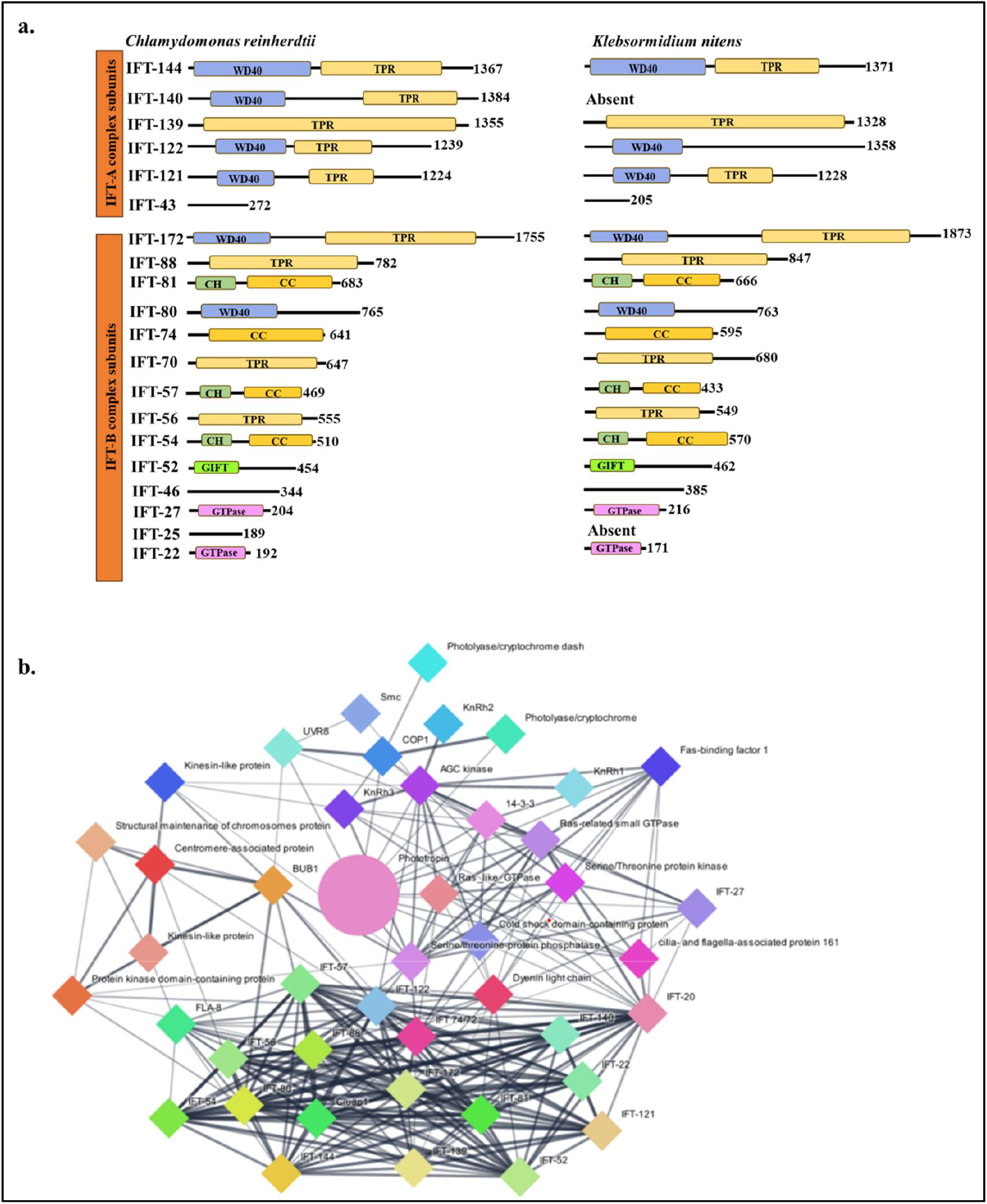
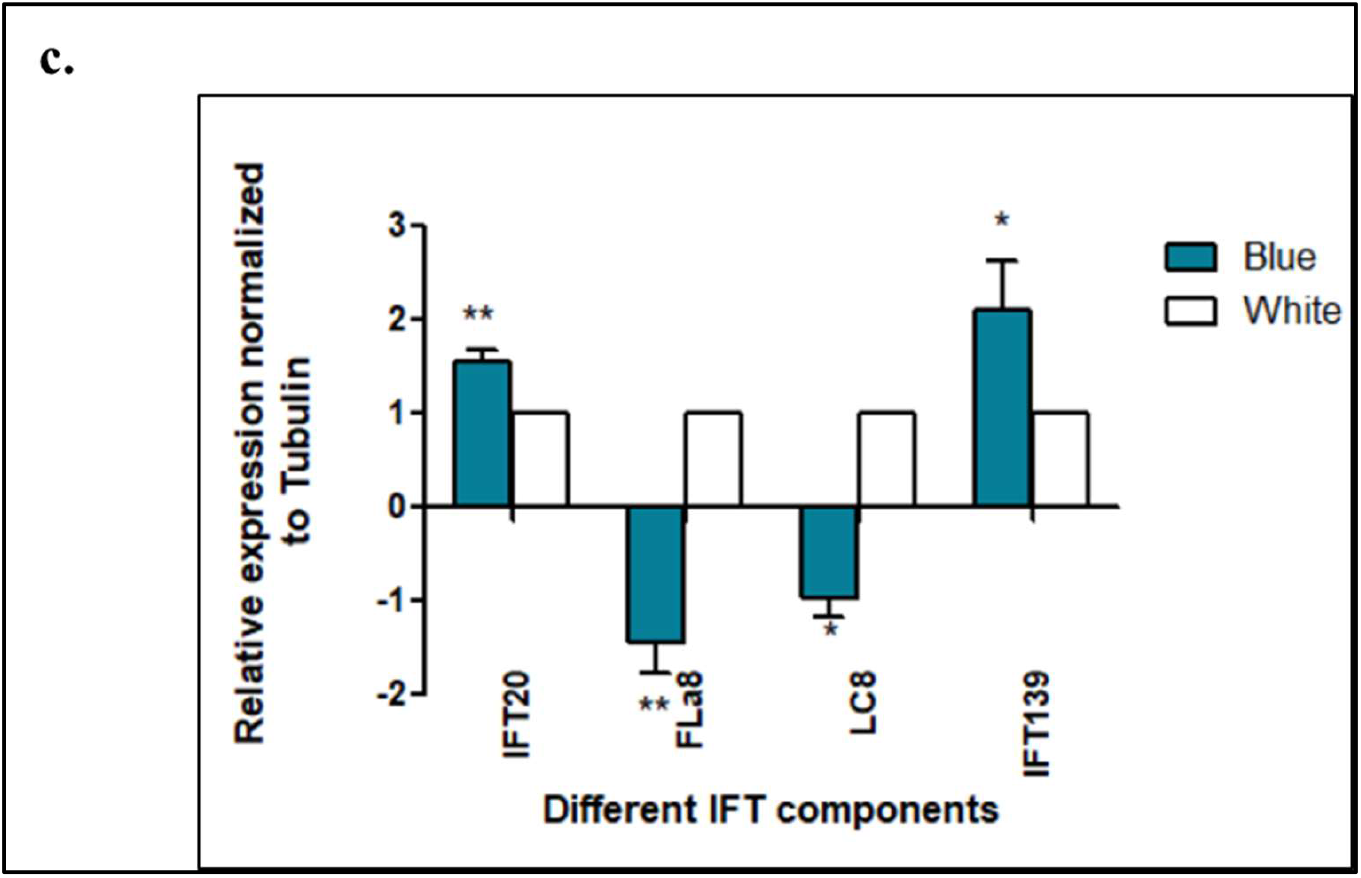
Schematic representation of intraflagellar transport (IFT) protein comparative domain architecture, interactome analysis and gene expression analysis of predicted protein interacting with phototropin in *Klebsormidium nitens*. a. Schematic domain representation of Intraflagellar transport components in *Chlamydomonas reinhardti* and *Klebsormidium nitens*. For the prediction of coiled coil (CC) https://toolkit.tuebingen.mpg.de/tools/pcoils tool was used. Tetratricopeptide repeats (TPR) were identified using https://toolkit.tuebingen.mpg.de/tools/tprpred and calponin homology (CH) domain were analysed using https://toolkit.tuebingen.mpg.de/tools/hhpred. Additional protein domains were identified through homology-based analysis and by comparing the with previously known protein structures. Conserved domain (CD) search tool (https://www.ncbi.nlm.nih.gov/Structure/cdd/wrpsb.cgi) was employed for validatation of domain predictions. b. Phototropin mediated protein–protein interaction network was constructed using STRING and visualized with Cytoscape. Phototropin is shown as the central node, interacting with numerous intraflagellar transport (IFT) components, including IFT20, IFT22, IFT27, IFT52, IFT57, IFT74/72, IFT81, IFT122, IFT139 and IFT172, as well as the motor proteins. The interactome highlights a potential functional association between phototropin and the intraflagellar transport machinery in *K. nitens*. c. Expression profile of different IFT components under white and blue light conditions in *Klebsormidium nitens*. Values are mean ± n=3, * and ** represents significance P<0.05 and P<0.01 respectively.

Notably, IFT140—a core IFT-A subunit that is essential for retrograde ciliary transport in *Chlamydomonas* and many other ciliated eukaryotes—is different from other organisms (*Chlamydomonas, Homo sapiens*), it differs in length as well as displays altered domain organisation. IFT122 is present, but it lacks the canonical TPR (tetratricopeptide repeat) motifs that are normally conserved in flagellated species and are required for stable IFT-A complex assembly. These structural alterations suggest a reduction or modification of classical IFT-A– mediated cargo transport functions/cytoskeleton organisation in *K. nitens*. IFT25-small IFT-B subunit, which forms a functional heterodimer with IFT27 and plays a key role in flagellar signalling in *Chlamydomonas*, is also absent. *K. nitens*, is filamentous and non-flagellated in the vegetative state. However, it has motile zoospore with inserted biflagella. The transition from vegetative state to motile zoospore state triggers by various environmental stimulus such as light, stress, and temperature shift. This suggests that the loss of IFT25–IFT27 module closely associated with flagellar membrane trafficking is dispensable in the non-flagellated alga, *K. nitens* in vegetative state.

As in *C. reinhardtii* the IFT acts as transport machinery for photoreceptors such as phototropin, we curated protein networking to understand the interaction of phototropin with IFT components. Protein-protein interaction network shows phototropin as the central hub protein within the network showing interaction with other photoreceptors (UVR8, KnRh1, KnRh2, Cryptochrome, Cryptochrome Dash) and IFT components such as IFT-139, IFT-20, IFT-88, IFT-52, IFT-70 (Sushmita et al., 2025) along with anterograde (kinesin) and retrograde (dynein) motor proteins (Fig. 2b, Table S2). The interaction of phototropin with other photoreceptors as well as with IFT-related transport components points to an integrated network that links signalling and intracellular trafficking. This connectivity supports the idea that IFT proteins contribute to the intracellular organization and spatial regulation of photoreceptor signalling in this non-flagellated vegetative state of alga.

Moreover, we also analysed the expression pattern of different representative genes of IFT components of each sub-complex (Fig. 2c). We found significant expression of the IFT component genes in the dark adapted cells. Additionally, we examined their expression pattern under blue light as compared to white light. The *IFT139* and *IFT20* gene expression was upregulated along with *phot* gene expression. Conversely, the motor proteins of each component were downregulated in blue light as compared to white light.

### Cellular detection of phototropin in *Klebsormidium nitens*

Following light activation, phototropins can dynamically relocalize, showing partial redistribution to the cytosol or to distinct membrane subdomains. The cellular localization of Phototropin has not yet explored in *K. nitens*. Thus, in the present study we performed the cellular localization of phototropin in *K. nitens* using lab raised antibody against LOV region of phototropin of *K. nitens*. The specificity of antibody was checked by immunoblotting with cell lysate. The presence of band of molecular weight approx. ∼85kDa shows specificity whereas, no band was observed in preimmune serum (Fig. 3a). Further, the immunofluorescence assay showed its localisation at the plasma membrane (Fig. 3b, S2).

**Fig. 3:**
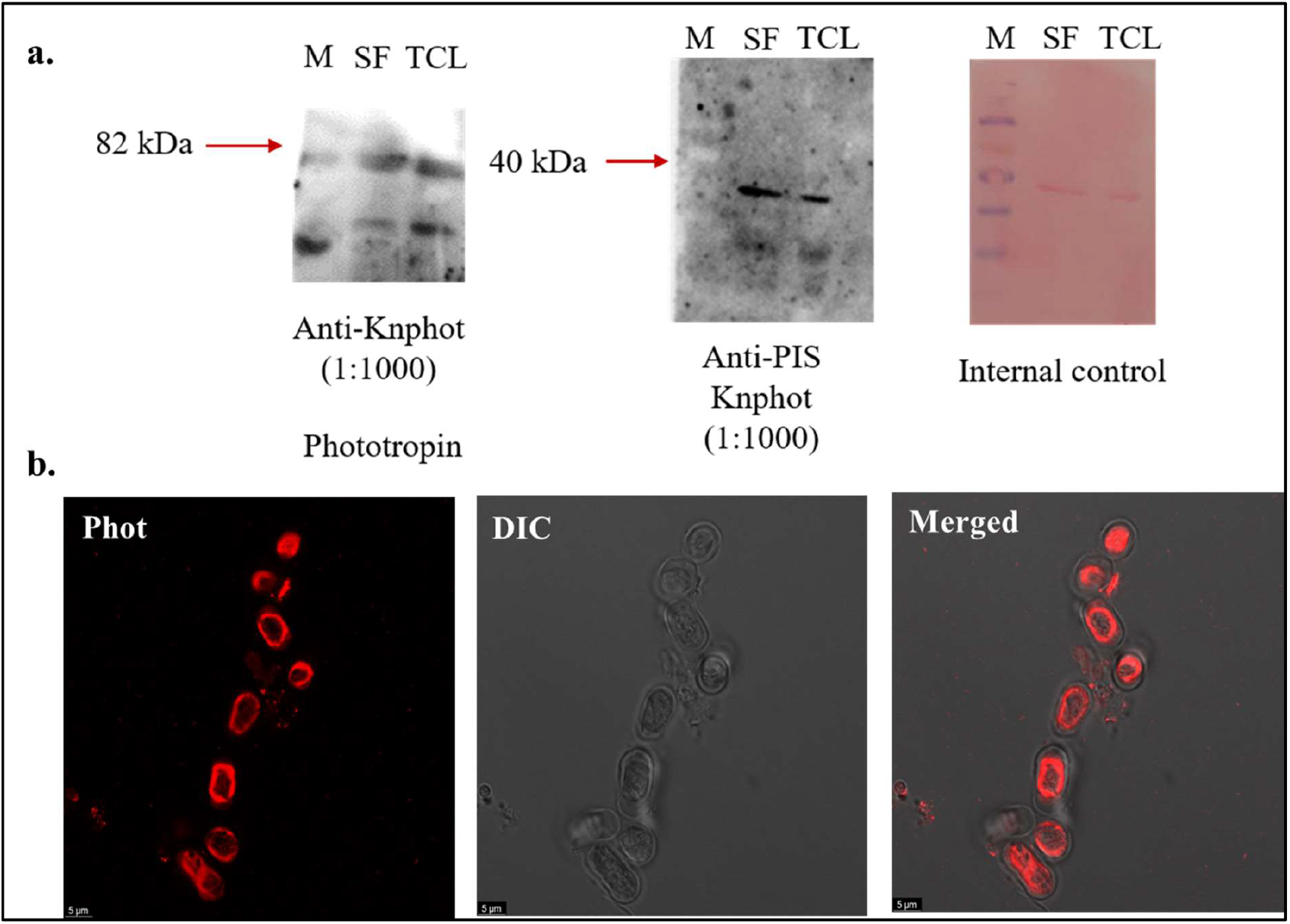
Cellular detection and localization of phototropin in *Klebsormidium nitens*. a. Immunoblot based cellular detection of *K*.*nitens* Phototropin. Total cell lysate (TCL) and Soluble fraction (SF) showing immunoreactive band around 85kDa probed with anti-Phot (dilution 1:1000) antibody indicated by red arrow. Pre-immune serum (PIS) used as a negative control. Ponceau-stained membranes are shown as a loading control. b. Immunolocalization of phototropin red channel using primary antibody against Knphot (dilution 1:1000) and secondary antibody anti-rabbit Alexa 546 (1:1000). Second channel represents DIC and third channel is the overlay of first two channels.

### Immunolocalization shows that phototropin spatial distribution might be linked with IFT-139

Intraflagellar transport protein 139 (IFT139) is a key part of the IFT-A complex. It functions as a peripheral subunit of this transport machinery and is best known for its role in powering retrograde transport, the process that moves materials from the tip of cilia and flagella back toward the cell body (Hirano et al., 2017; Nishat et al., 2025). As observed in Fig. 2b, *K. nitens* phototropin was found to interact with IFT-139. Immunoblotting and immunofluorescence assay suggest the presence of IFT139 in *K. nitens* (Fig. 4). Immunoblotting detection was done using lab raised antibody against CrIFT-139 antigen from *C. reinhardtii*. Our results showed a clear band at ∼130kDa in both total cell lysates and soluble fraction of *K. nitens*. In contrast, no corresponding signal was detected in the control lanes, demonstrating the specificity of the antibody. However, we also found bands of different molecular weight (Fig. 4a).

**Fig. 4:**
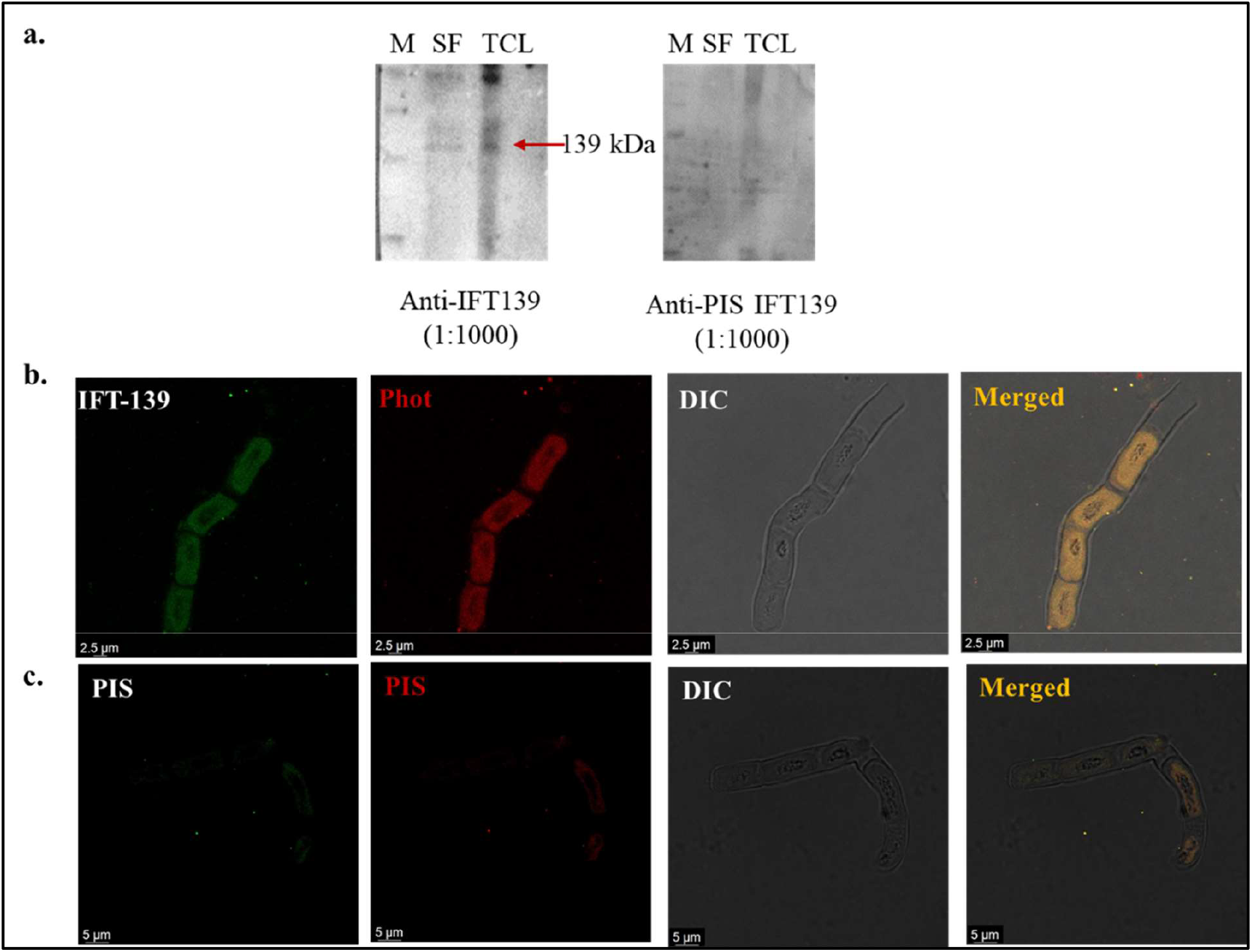
Cellular detection and immunolocalization of the IFT-A complex component, IFT-139 colocalization with phototropin in *Klebsormidium nitens*. a. Immunoblot analysis confirmed the presence of IFT-139, a core component of the IFT-A complex, in *K. nitens*. Total cell lysate (TCL) and soluble fraction (SF) probed with anti–IFT-139 antibody (dilution1:1000) shows a band around 130kDa marked by the red arrow, verifying the expression of the IFT-139 protein. b. Immunofluorescence microscopy revealed the cellular distribution of IFT-139 (green) and phototropin (Phot; red) in *K. nitens*. Green channel (first panel) indicates the detection of IFT-139 using anti-CrIFT-139 as primary antibody (1:250) and secondary antibody Alexa Fluor 488-conjugated anti-goat antibody. Red channel (second panel) shows the detection of KnPhot using anti-KnPhot as primary antibody (1:1000) and Alexa Fluor 546-conjugated anti-rabbit secondary antibody. Differential interference contrast (DIC) images shown in panel third provide structural context, while fourth panel shows merged images indicates spatial overlap between IFT-139 and phototropin signals. Scale bar= 2.5µm. c. First panel shows pre-immune serum using primary anti-PISCr139 (dilution-1:250) and secondary antibody anti-goat IgG Alexa-488 (dilution-1:1000). Second panel indicates pre-immune serum using primary anti-KnPhot (dilution-1:1000) and secondary antibody Alexa Fluor 546-conjugated anti-rabbit antibody. Third panel shows DIC image of *K*.*nitens* cells. Fourth panel indicates the overlay images of all three panels. Scale bar= 5µm.

Furthermore, immunofluorescence microscopy revealed a distinct intracellular distribution along of IFT139 along the length (cortical regions) of filamentous cells of *K. nitens*. Interestingly, our co-localization analysis demonstrated substantial overlap between the IFT-139 (green) and phototropin (red) signals in merged images (Fig. 4b, S3), indicating a close spatial association between these two proteins. No detectable signal was observed in the pre-immune serum controls, further confirming the specificity of the observed localization pattern (Fig. 4c). Importantly, this interaction and co-localization identified in a filamentous non-flagellated vegetative state of alga, suggests that IFT-139 has been retained and functionally repurposed beyond its canonical role in flagellar transport. Unlike *C. reinhardtii*, where IFT-139 is an essential component of the flagellar IFT-A complex, the distribution pattern observed in *K. nitens* supports a non-ciliary function for IFT-139, potentially related to phototropin trafficking or the spatial organization of light-signalling components (Sanyal et al., 2023a; Sanyal et al., 2023b; Sharma et al., 2024).

### IFT 20 shows co-localization with phototropin along the cortical length

Intraflagellar transport protein 20 (IFT-20) is a highly conserved part of the IFT-B complex and is crucial for anterograde intraflagellar transport. This process is essential for building, maintaining, and ensuring the proper function of cilia and flagella in eukaryotic cells. IFT 20 is known for its dual localization: in addition to being present in cilia, it is also found at the Golgi apparatus and within cytoplasmic vesicle trafficking pathways. In flagellated organisms such as *Chlamydomonas reinhardtii*, IFT-20 plays a key role in transporting membrane and signaling proteins to the cilium, effectively linking intracellular vesicle trafficking with microtubule-based transport along the axoneme (Follit et al., 2006; Pazour & Bloodgood, 2008).To confirm the presence of IFT-20 *in K. nitens* immunoblotting was performed using anti-CrIFT-20 antibody. Immunoblot analysis confirmed the IFT-20 subunit protein in total cell lysate as well as soluble fraction with immunoreactive band detected ∼ 20 kDa. No signal was detected in the control lanes, further validating antibody specificity (Fig. 5a). Further, immunolocalization study revealed that it is co-localized with phototropin suggesting that spatial distribution of phototropin might be linked with IFT20 in *K. nitens* (Fig. 5b, S4). IFT-20 showed strong localization at the periphery of the filamentous cells and cortical region. Phototropin displayed a closely matching pattern, with predominant accumulation along the cortical region of the cell. Merged fluorescence imaging showed a strong overlap between IFT-20 (green) and phototropin (red), producing clear yellow signals along the filaments and indicating robust co-localization of the two proteins. Differential interference contrast (DIC) images verified that these fluorescence signals originated from intact filamentous cells. By contrast, samples treated with pre-immune serum displayed no detectable fluorescence, confirming that the observed localization patterns were specific.

**Fig. 5:**
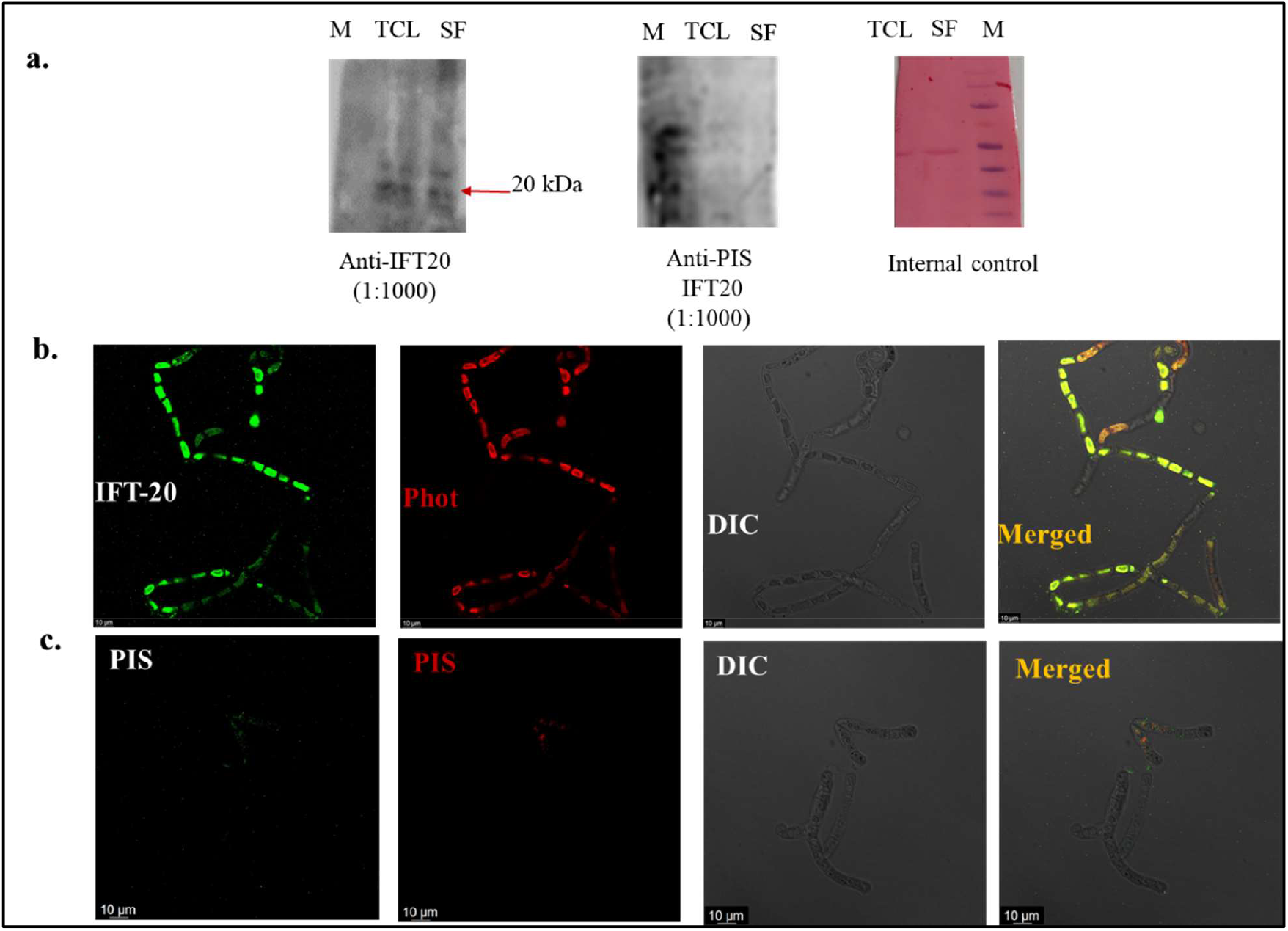
Cellular detection and immunolocalization of the IFT-B complex component IFT-20 and its colocalization with *K. nitens* phototropin (Phot). a. Immunoblot analysis showing detection of IFT-20 in *K*.*nitens* in total cell lysate (TCL) and soluble fraction (SF) probed with *Chlamydomonas* IFT20 (CrIFT20) antibody (1:1000) shows a band around 20kDa. Negative signal in pre-immune serum shows antibody specificity. b. Immunofluorescence microscopy showing the cellular distribution of IFT-20 (green) and phototropin (Phot; red) in *K. nitens*. In first panel, IFT-20 was detected using anti-CrIFT-20 as the primary antibody (1:250) followed by anti-goat Alexa Fluor 488-conjugated secondary antibody (1:1000 dilutuon), while the second panel *K. nitens* phototropin was detected using anti-Phot primary antibody (1:1000) followed by anti-rabbit Alexa Fluor 546-conjugated secondary antibody. The third panel shows differential interference contrast (DIC) images that provide cellular morphology, and the fourth panel corresponds to merged images that reveal colocalization of IFT-20 and phototropin. Scale bar = 10 µm. c. The first panel shows immunolocalization with pre-immune serum for CrIFT-20 and KnPhot (dilution-1:1000) and secondary antibody anti-goat IgG Alexa-488 (dilution-1:1000). Second panel indicates pre-immune serum using primary anti-KnPhot (dilution-1:1000) and secondary antibody Alexa Fluor 546-conjugated anti-rabbit antibody. The third panel shows DIC image of *K. nitens* cells. The fourth panel indicates the overlay images of all three panels. Scale bar= 10µm.

Taken together, these findings demonstrate a close association between phototropin and the IFT-20 subunit in *K. nitens*. The cortical co-localization of phototropin with an intraflagellar transport component in this filamentous alga supports the idea that IFT machinery has been evolutionarily repurposed for non-ciliary roles, potentially facilitating the spatial organization or intracellular trafficking of phototropin-dependent light-signaling components along microtubule-associated cortical regions.

### The motor protein of anterograde and retrograde IFT subcomplexes shows similar spatial distribution as phototropin in *K. nitens*

The motor protein LC8, dynein light chain, and Fla8, kinesin-2 motor protein, are motor proteins for retrograde (IFTA) and anterograde (IFTB) sub-complexes. FLA8 is a subunit of the kinesin-II motor complex that powers anterograde, microtubule-based transport during intraflagellar transport (IFT) (Pedersen et al., 2006). In contrast, LC8 (dynein light chain 8) is a highly conserved accessory protein that binds to dynein motors and acts as a flexible dimerization hub, helping connect cargo to the cytoskeleton (Barbar, 2008; Rapali et al., 2011). In our investigation, we found the presence of LC8 and Fla8 via cellular detection using lab raised antibody generated against anti-LC8 and anti-Fla8 antigen from *C. reinhardtii* (Fig. 6 and 7). We observed the spatial distribution of LC8 and Fla8 was same as phototropin. They showed colocalization with phototropin along the cortical region (Fig. 6b and 7b, S5 and 6). This suggests that LC8 and Fla8 are present in *K. nitens*, but serve cytoplasmic roles potentially linking to phototropin trafficking. No signal was observed in pre immune serum (Fig. 6c and 7c). The detection of two distinct molecular forms of LC8 and FLA8 likely points to post-translational control mechanisms acting on motor proteins and components associated with intraflagellar transport (IFT). Both the dynein light chain LC8 and the kinesin-II subunit FLA8 have established roles in regulated transport pathways, where phosphorylation-dependent shifts in electrophoretic mobility have previously been observed for motors and their adaptors. These modifications are thought to fine-tune motor function by influencing activity, cargo interactions, or engagement with IFT particles, aligning well with the highly dynamic nature of intracellular trafficking mediated by these proteins (DeBerg et al., 2013; Dillman & Pfister, 1994).The idea that conserved IFT and motor components are still under regulatory control even in non-flagellated lineages is further supported by the biochemical heterogeneity of LC8 and FLA8 seen in *K. nitens*.

**Fig. 6:**
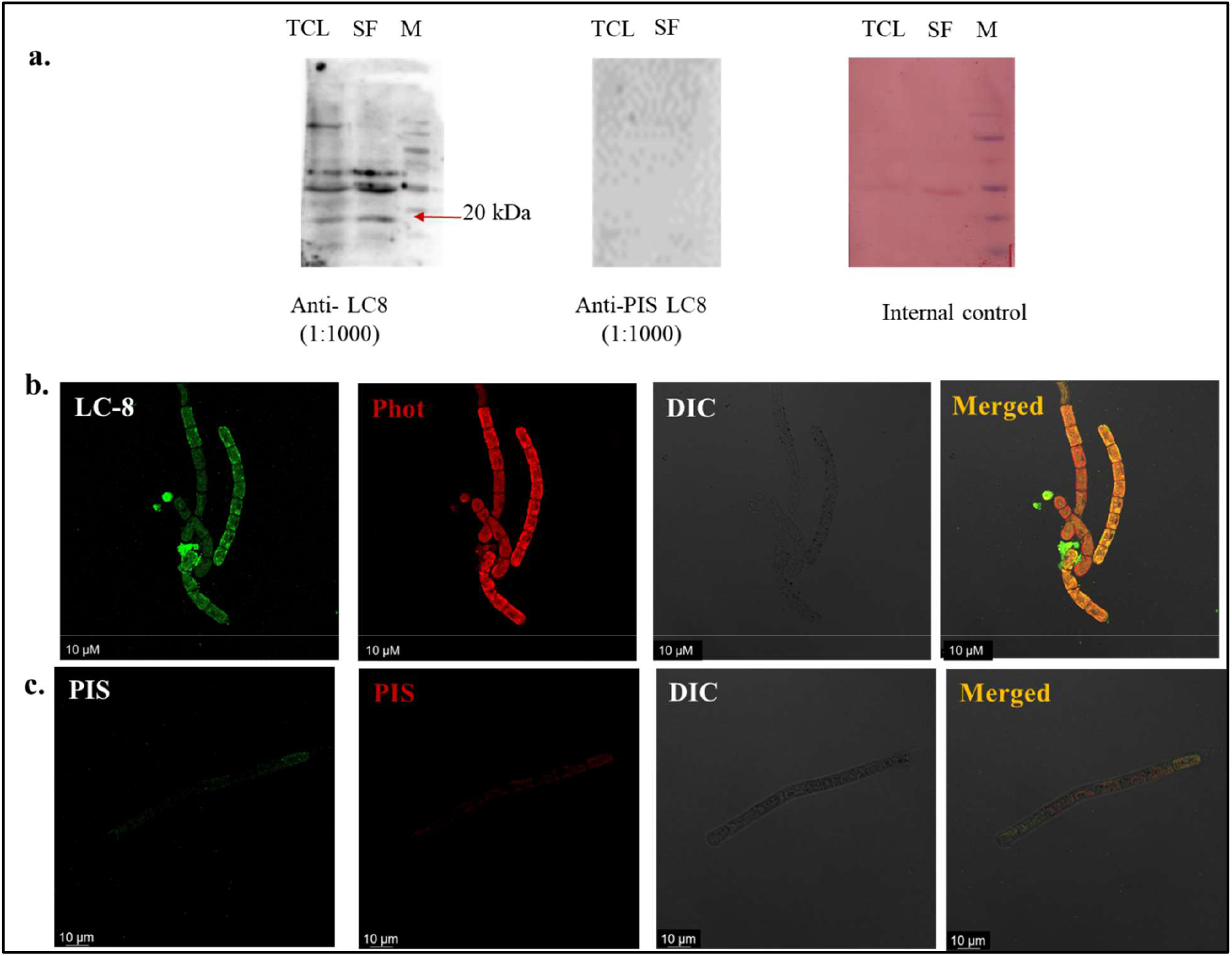
Detection and cellular localization of the anterograde motor protein LC-8 and its colocalization with phototropin in *Klebsormidium nitens*. a. Immunoblot analysis showing detection of LC-8 in *K*.*nitens* in total cell lysate (TCL) and soluble fraction (SF) probed with CrLC8 antibody (1:1000) shows a band at ∼20kDa. Negative signal in pre-immune serum shows antibody specificity. Ponceau serving as an internal control. b. Immunofluorescence microscopy showing the cellular distribution of LC8 (green) and phototropin (Phot; red) in *K. nitens*. In the first panel LC8 was detected using anti-CrLC8 as the primary antibody (1:250), followed by Alexa Fluor 488-conjugated secondary antibody, while in the second panel *K. nitens* phototropin was detected using anti-Phot primary antibody (1:1000), followed by Alexa Fluor 546-conjugated secondary antibody. The third panel shows differential interference contrast (DIC) images that provide cellular morphology, and the fourth panel corresponds to merged images that reveal colocalization of LC8 and phototropin. Scale bar = 10 µm. c. The first panel shows pre-immune serum using primary pre-immune serum for primary CrLC8 antibody (dilution-1:250) and secondary antibody anti-goat IgG Alexa-488 (dilution-1:1000). The second panel indicates pre-immune serum using primary anti-KnPhot (dilution-1:1000) and secondary antibody Alexa Fluor 546-conjugated anti-rabbit antibody. The third panel shows DIC image of *K. nitens* cells. The fourth panel indicates the overlay images of all three panels. Scale bar= 10µm.

**Fig. 7:**
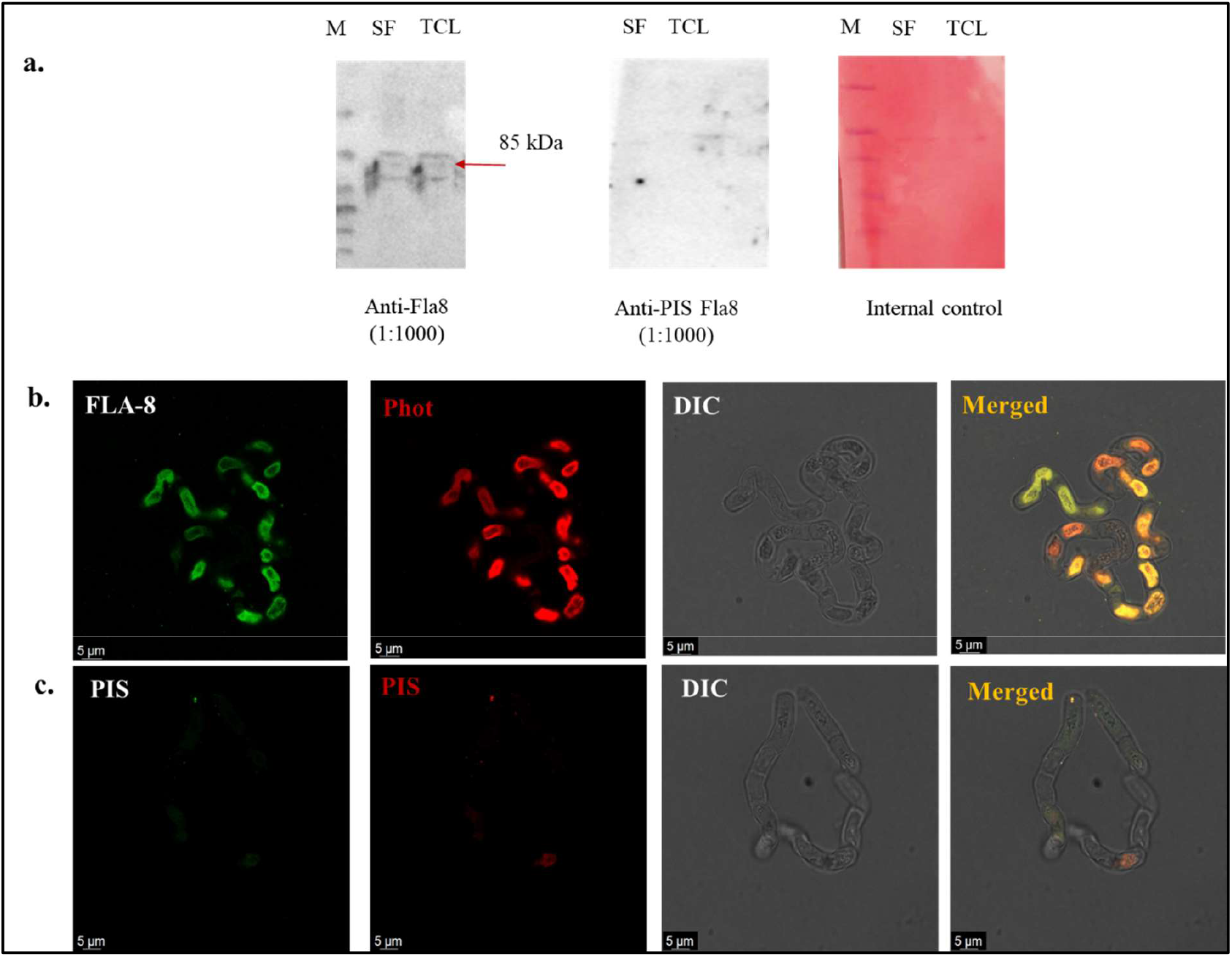
Cellular detection and immunolocalization of the retrograde motor protein FLA-8 and its colocalization with *K. nitens* phototropin (Phot). a. Immunoblot analysis showing detection of FLA-8 in *K*.*nitens* in total cell lysate (TCL) and soluble fraction (SF) probed with CrFLA8 antibody (1:1000) shows a band ∼85kDa. No signal in pre-immune serum shows antibody specificity. Ponceau stain was used as an internal control to show equal loading. b. Immunofluorescence microscopy showing the cellular distribution of FLA-8 (green) and phototropin (Phot; red) in *K. nitens*. In the first panel, FLA-8 was detected using anti-CrFLA8 as the primary antibody (1:250) followed by Alexa Fluor 488-conjugated secondary antibody, while in the second panel *K. nitens* phototropin was detected using anti-KnPhot primary antibody (1:1000), followed by anti-rabbit Alexa Fluor 546-conjugated secondary antibody. The third panel shows differential interference contrast (DIC) images that shows cellular morphology, and the fourth panel corresponds to merged images that reveal colocalization of FLA-8 with phototropin. Scale bar = 5 µm. c. First panel shows pre-immune serum FLA-8 using pre-immune serum for primary CrFLA8 antibody (dilution-1:250) and secondary antibody anti-goat IgG Alexa-488 (dilution-1:1000). The second panel indicates pre-immune serum using primary anti-KnPhot (dilution-1:1000) and secondary antibody Alexa Fluor 546-conjugated anti-rabbit antibody (1:1000). The third panel shows DIC image of *K*.*nitens* cells. The fourth panel indicates the overlay images of all three panels. Scale bar= 10µm.

### *Klebsormidium nitens* 14-3-3 association with phototropin is found to be conserved

14-3-3 proteins are highly conserved regulatory factors which exist either in homodimers or heterodimers. They act as adaptor or scaffold proteins by binding to phosphorylated target proteins and influencing their stability, localization, and activity (Fu et al., 2000; Van Heusden, 2005). In land plants, 14-3-3 proteins are well-known components of phototropin signalling pathways, where they interact directly with phototropin or with phototropin-regulated proteins to drive downstream responses such as stomatal opening and ion transport (Inoue & Kinoshita, 2017; Reuter et al., 2021; Tseng et al., 2012). It is widely conserved across eukaryotic lineages and maintain key structural and functional characteristics, including conserved phosphorylation, dimerization interfaces, and calcium-binding motifs that are essential for their activity. In line with this evolutionary conservation, in the present study we performed the sequence alignment analysis of 14-3-3 with higher organisms (Fig. S7). It showed that the *K. nitens* 14-3-3 protein shares a high level of sequence similarity with 14-3-3 homologs from higher plants and *Chlamydomonas reinhardtii*. Notably, residues required for phosphorylation, dimer formation, and regulatory function are strongly conserved, suggesting that the *K. nitens* 14-3-3 protein is functionally active. Immunofluorescence analysis further revealed that phototropin exhibits a distinct intracellular distribution along the filamentous cells of *K. nitens*, with strong association at or near the plasma membrane. This pattern closely resembles/mirrors the plasma membrane–associated localization of phototropins observed in higher plants, supporting the idea that plant-like spatial regulation of phototropins arose prior to land colonization. Notably, 14-3-3 proteins displayed a similar intracellular distribution and showed clear colocalization with phototropin (Fig. 8). The specificity of lab raised antibody against 14-3-3 of *C. reinhardtii* was checked with preimmune serum using immunoblotting and immunofluorescence (Fig. 8a and c). The overlapping fluorescence signals in merged images indicate that the phototropin–14-3-3 interaction in *K. nitens* occurs independently of flagellar structures (Fig. 8b, S8). This points at fundamentally different spatial organization in *C. reinhardtii*, where phototropin localization is tightly linked to flagella. The non-flagellar, plasma membrane–associated colocalization of phototropin and 14-3-3 proteins in *K. nitens* therefore represents a fundamentally different mode of blue-light signaling, one that more closely resembles the phototropin signalling architecture characteristic of land plants.

**Fig. 8:**
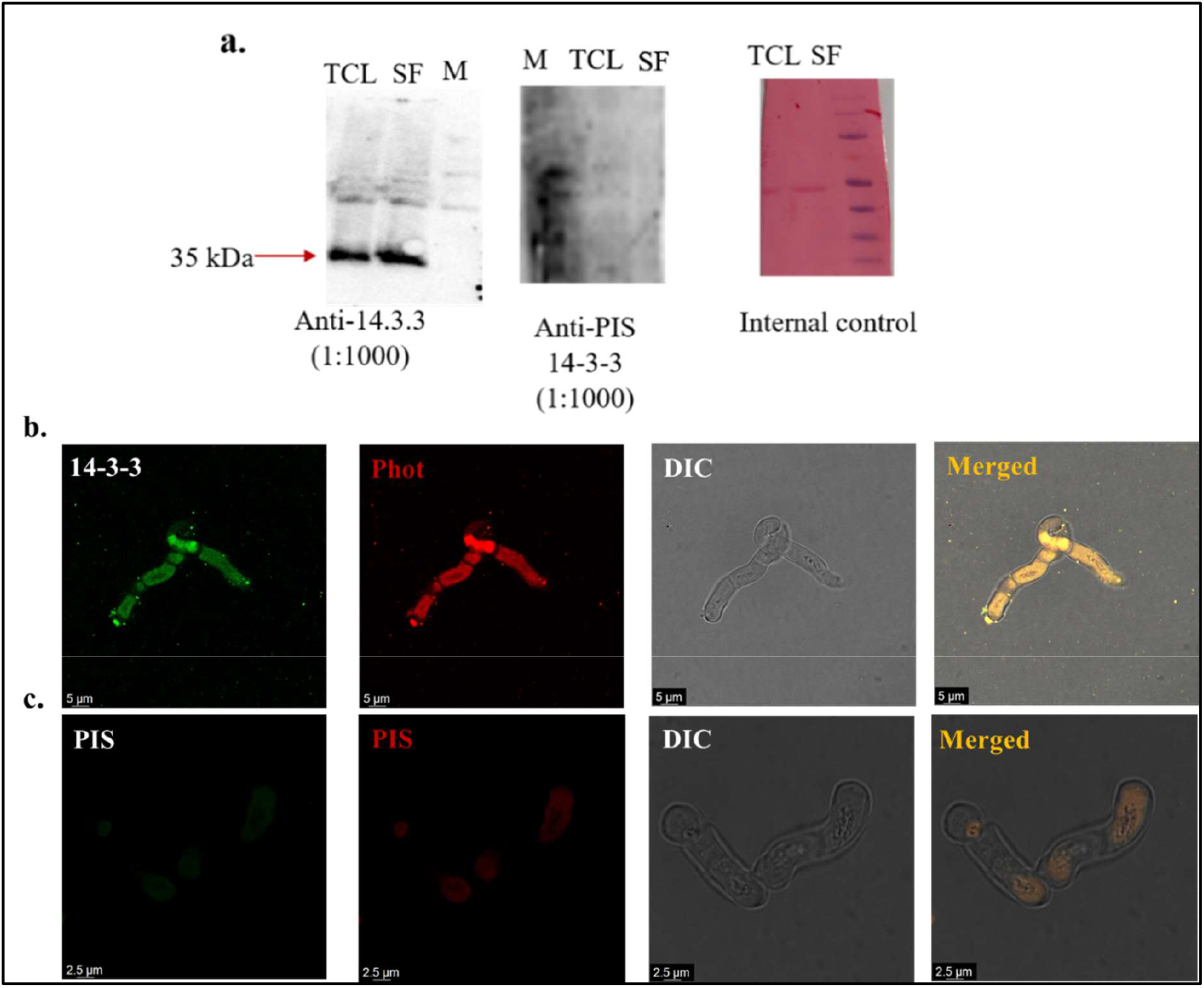
Cellular localisation of 14-3-3 and its co-localisation with phototropin in *Klebsormidium nitens*. a. Immunoblot based cellular detection of *K. nitens* 14-3-3. Total cell lysate (TCL) and Soluble fraction (SF) showing immunoreactive band around 35kDa probed with anti-Cr14-3-3 antibody indicated by red arrow confirms the presence of phototropin in *K. nitens*. Absence of signal with preimmune serum confirms signal specificity for presence of protein. Ponceau stained membranes are shown as loading control. b. Cellular localization of 14-3-3 protein and phototropin in *K. nitens* using immunofluorescence microscopy. In the upper panel, the green channel (14-3-3) shows 14-3-3 protein using primary anti-Cr14-3-3 antibody (dilution 1:250) and Alexa Fluor 488-conjugated anti-goat secondary antibody. The red channel (KnPhot) localisation was detected by using anti-KnPhot primary antibody (1:1000) followed by Alexa Fluor 546-conjugated anti-rabbit secondary antibody. The third panel shows that differential interference contrast (DIC) is used to show cellular morphology. The fourth panel shows merged images indicating co-localisation of 14-3-3and phototropin. Scale bar=5µm. c. Lower panels show pre-immune serum (PIS) 14-3-3 using anti-PIS Cr14-3-3(1:250), followed by anti-goat Alexa Fluor 488-conjugated secondary antibody and PIS KnPhot using anti-PIS KnPhot (1:1000), followed by anti-rabbit Alexa-Fluor 546 -conjugated secondary antibody, representing controls for both antibodies, confirming signal specificity. Scale bars = 2.5 µm.

The observed colocalization pattern of phototropins with 14-3-3 protein in *K. nitens* points to a fascinating evolutionary repurposing of intracellular machinery. Traditionally recognized as adaptors for phosphorylated intraflagellar transport (IFT) and motor-associated proteins, 14-3-3 proteins appear here to operate at the plasma membrane, suggesting that ancestral IFT-linked signaling modules have been co-opted to regulate non-ciliary light responses. This finding provides compelling evidence that early plant lineages retained and adapted preexisting ciliary signaling pathways to meet the demands of light perception and signaling outside of cilia (Kumari et al., 2025; Kumari & Ray, 2022; Sharma et al., 2025).

## Conclusions

This study shows that key parts of the intraflagellar transport (IFT) system have been preserved and reassigned a new functions in the vegetative state of filamentous streptophyte alga *K. nitens*. Although this organism has completely lost motile cilia and flagella, majority of IFT-A and IFT-B subunits are still present, indicating presence of IFT machinery before flagellar loss. Thus, indicating evolutionarily important role in cellular functions unrelated to cilia. The co-localization of IFT components with phototropin suggests their localization is cell cortex and plasma membrane-associated in *K. nitens*. This pattern is strikingly different from the classical axonemal and basal body localization seen in flagellated algae like *C. reinhardtii*, but closely resembles the membrane-associated distribution of phototropin observed in land plants. These observations suggest that, in *K. nitens*, IFT proteins might be involved in membrane-based trafficking or the spatial organization of signaling complexes rather than axonemal transport. . Given the known role of 14-3-3 proteins in regulating cargo binding and motor interactions, our findings hypothesized a mechanism in which 14-3-3 acts as a molecular bridge linking phototropin to IFT-derived trafficking modules outside of a ciliary context. Together, these results demonstrate that the ancestral IFT machinery has been repurposed to support photoreceptor organization and blue-light signaling in a filamentous, non-flagellated alga in vegetative state. This functional shift likely represents an evolutionary transition between the flagellum-based signaling systems of unicellular algae and the membrane-based photoreceptor signaling pathways found in land plants. Overall, our work highlights the evolutionary flexibility of the IFT system and underscores its broader role in intracellular signaling beyond ciliary transport.

## Data Statement

Data will be made available on request.

## Author’s approval

All authors have seen and approved the manuscript. This work is original and not under consideration or published anywhere.

## Author Contributions

S.K. conceived the project. M. and R.S. designed, performed the experiments and drafted the manuscript in the guidance of S.K. S.S. generated the *Klebsormidium nitens* phototropin antibody and edited manuscript. All authors reviewed and approved the final manuscript.

## Acknowledgements

SK is grateful to ANRF/SERB for providing research grants (EEQ/2023/000398). M. and S.S. acknowledge CSIR-UGC for providing the Junior research fellowship and Senior research fellowship (NTA Ref. No. 211610173246). RS acknowledge DBT-RA program (DBT-RA/2022/July/N/2560).

