## Supplementary materials for "Ancient intraflagellar transport machinery controls unique spatial distribution of phototropin in an evolutionary important non-flagellated vegetative stage of terrestrial alga"

^#^ Current Affiliation

* To whom correspondence should be addressed.

†ORCID identifiers: M. 0009-0001-7374-1788, R.S. 0000-0002-1520-2794, S.S. 0000-0002-6531-7522, S.K. 0000-0001-5428-4297

**Supplementary Tables**

**Table S1: List of primers used for qRT-PCR analysis in this study.**

| ***Primer*** | ***Sequence (5’-3’)*** | **References** |
| --- | --- | --- |
| **Phototropin** | CTCTCTCCACGTTCAAGCATAC  CTGGTCCATATTGCGTCATCTC | This study |
| **IFT-139** | CGTGGATCGAGTTGTCATCA  CCCAACACCTCCTCAAAGAA | This study |
| **IFT-20** | GCAGAGAAGCAACAGGAACTA  CGTCTGTGATCTTCGCTATCAT | This study |
| **FLA-8** | GATGCGGAAGGAAATGGAGA  CTCCCGCTCTTTCATCATACTC | This study |
| **LC-8** | CCTGAGAAGACCAGAGATGAA  CAAATTCCCTGGCCATAGTG | This study |

**Table S2: List of identified Intraflagellar transport components in *Klebsormidium nitens* in reference to *Chlamydomonas reinhardti* along with their accession number.**

| **IFT components** | **Uniport ID (*C. reinhardtii*)** | **Accession number (*K. nitens*)** |
| --- | --- | --- |
| **IFT-144** | **A9XPA7** | **GAQ81709** |
| **IFT-140** | **Q68K27** | **GAQ77910** |
| **IFT-139** | **A9XPA6** | **GAQ89264** |
| **IFT-122** | **H9CTG6** | **GAQ89710** |
| **IFT-121** | **A8JFR3** | **GAQ80981** |
| **IFT-43** | **A8HYP5** | **GAQ80924** |
| **IFT-172** | **Q5DM57** | **GAQ82168** |
| **IFT-81** | **Q68RJ5** | **GAQ81647** |
| **IFT-88** | **A8JCJ2** | **GAQ81294** |
| **IFT-80** | **A5Z0S9** | **GAQ90821** |
| **IFT-74** | **Q6RCE1** | **GAQ77667** |
| **IFT-70** | **A8ITN7** | **GAQ82289** |
| **IFT-57** | **Q2XQY7** | **GAQ80546** |
| **IFT-56** | **A8JA42** | **GAQ81671** |
| **IFT-54** | **A8JBY2** | **GAQ86389** |
| **IFT-52** | **Q944U2** | **GAQ81859** |
| **IFT-46** | **A2T2X4** | **GAQ84005** |
| **IFT-38** | **A0A2K3CQB1** | **GAQ92317** |
| **IFT-27** | **A8HN58** | **GAQ83747** |
| **IFT-25** | **B8LIX8** | **Absent** |
| **IFT-22** | **A8HME3** | **GAQ79275** |
| **IFT-20** | **Q8LLV9** | **GAQ82347** |

**Table S3: List of different nodes and their Uniport ID for the interacting partners of phototropin and IFT components.**

| Node | Uniport ID |
| --- | --- |
| Ras_like_GTPase | **A0A0U9HIK3** |
| Dyenin light chain | **A0A1Y1HRE9** |
| IFT-20 | **A0A1Y1I0N5** |
| IFT 74/72 | **A0A1Y1HM71** |
| Phototropin | **A0A1Y1HNG4** |
| cilia- and flagella-associated protein 161/ EF-hand domain-containing protein | **A0A1Y1ILL8** |
| 14-3-3 | **A0A0U9HJD4** |
| Serine/Threonine protein kinase | **A0A1Y1HNN5** |
| Serine/threonine-protein phosphatase | **A0A1Y1HWR5** |
| AGC kinase ((cAMP-dependent, cGMP-dependent and protein kinase C) | **A0A0U9HKE2** |
| Ras-related small GTPase | **A0A1Y1IHY9** |
| KnRh3 | **A0A1Y1HT90** |
| IFT-27 | **A0A1Y1I4U9** |
| Fas-binding factor 1 | **A0A1Y1HIG9** |
| Cold shock domain-containing protein | **A0A1Y1I7V9** |
| Kinesin-like protein | **A0A1Y1HYT8** |
| Smc | **A0A1Y1I1A7** |
| COP1 | **A0A1Y1HGF7** |
| IFT-122 | **A0A1Y1IFT9** |
| KnRh2 | **A0A1Y1IBK3** |
| KnRh1 | **A0A1Y1I5V9** |
| Photolyase/cryptochrome dash | **A0A1Y1HMX6** |
| UVR8 | **A0A1Y1IBI5** |
| Photolyase/cryptochrome | **A0A1Y1IHG3** |
| IFT-140 | **A0A1Y1HHG8** |
| FLA-8 | **A0A1Y1IPN0** |
| IFT-22 | **A0A1Y1HKZ1** |
| Cluap1 | **A0A1Y1IUG9** |
| IFT-57 | **A0A1Y1HVJ0** |
| IFT-81 | **A0A1Y1HYM9** |
| IFT-56 | **A0A1Y1HXI3** |
| IFT-54 | **A0A1Y1IAD1** |
| IFT-52 | **A0A1Y1HXF8** |
| IFT-88 | **A0A1Y1HRN1** |
| IFT-172 | **A0A1Y1I21** |
| IFT-80 | **A0A1Y1INS1** |
| IFT-139 | **A0A1Y1IE71** |
| IFT-144 | **A0A1Y1HX18** |
| IFT-121 | **A0A1Y1HVH3** |
| Structural maintenance of chromosomes protein | **A0A1Y1IGP6** |
| BUB1 | **A0A1Y1HKE3** |
| Protein kinase domain-containing protein | **A0A1Y1IG22** |
| Kinesin-like protein | **A0A1Y1IH75** |
| Centromere-associated protein/ Kinetochore protein NDC80 | **A0A1Y1ILS8** |

**Supplementary Figures**

**Figure S1**

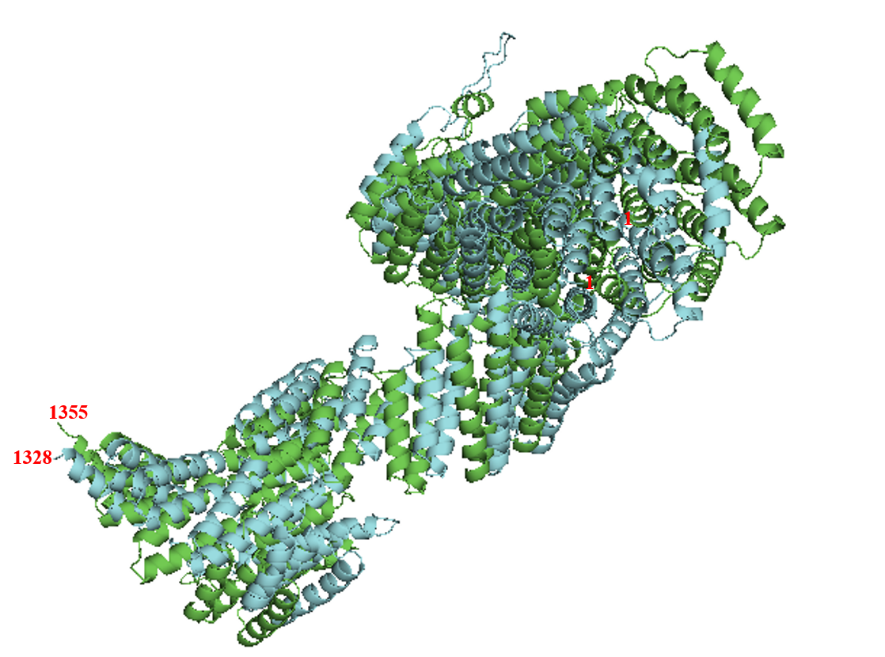

Fig. S1a: Superimposition of *Chlamydomonas reinhardtii* and *Klebsormidium nitens* IFT-139 structure. Structures were predicted using Alpha fold server <https://alphafoldserver.com/> and visualised using Pymol software. Green colour represents the *C. reinherdtii* IFT-139, whereas blue colour represents *K. nitens* IFT-139.

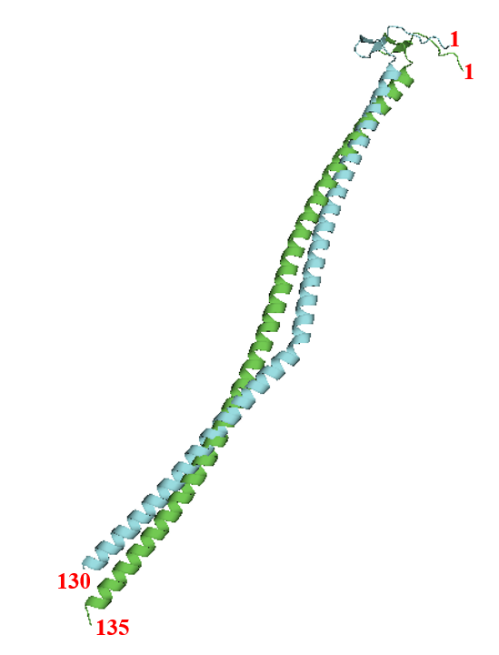

Fig. S2b: Superimposition of *Chlamydomonas reinhardtii* and *Klebsormidium nitens* IFT-20 structure. Structures were predicted using Alpha fold server <https://alphafoldserver.com/> and visualised using Pymol software. Green colour represents the *C. reinherdtii* IFT-20, whereas blue colour represents *K. nitens* IFT-20.

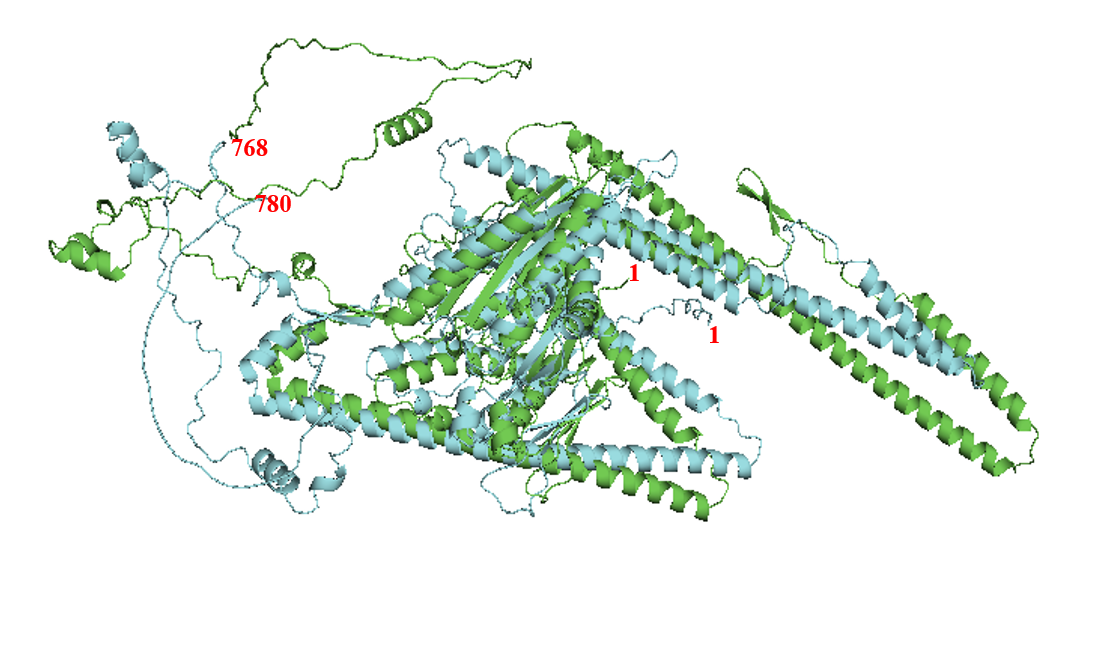

Fig. S1c: Superimposition of *Chlamydomonas reinhardtii* and *Klebsormidium nitens* FLA-8 structure. Structures were predicted using Alpha fold server <https://alphafoldserver.com/> and visualised using Pymol software. Green colour represents the *C. reinherdtii* FLA-8, whereas blue colour represents *K. nitens* FLA-8.

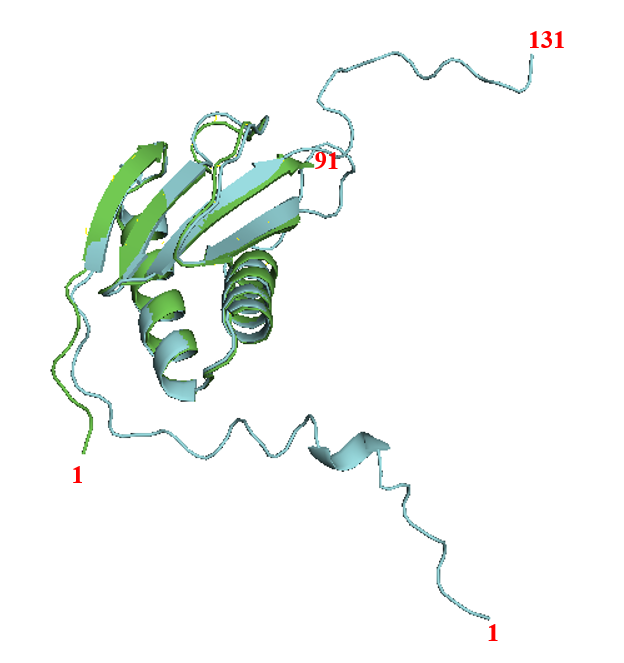

Fig. S1d: Superimposition of *Chlamydomonas reinhardtii* and *Klebsormidium nitens* LC-8 structure. Structures were predicted using Alpha fold server <https://alphafoldserver.com/> and visualised using Pymol software. Green colour represents the *C. reinherdtii* LC-8, whereas blue colour represents *K. nitens* LC-8

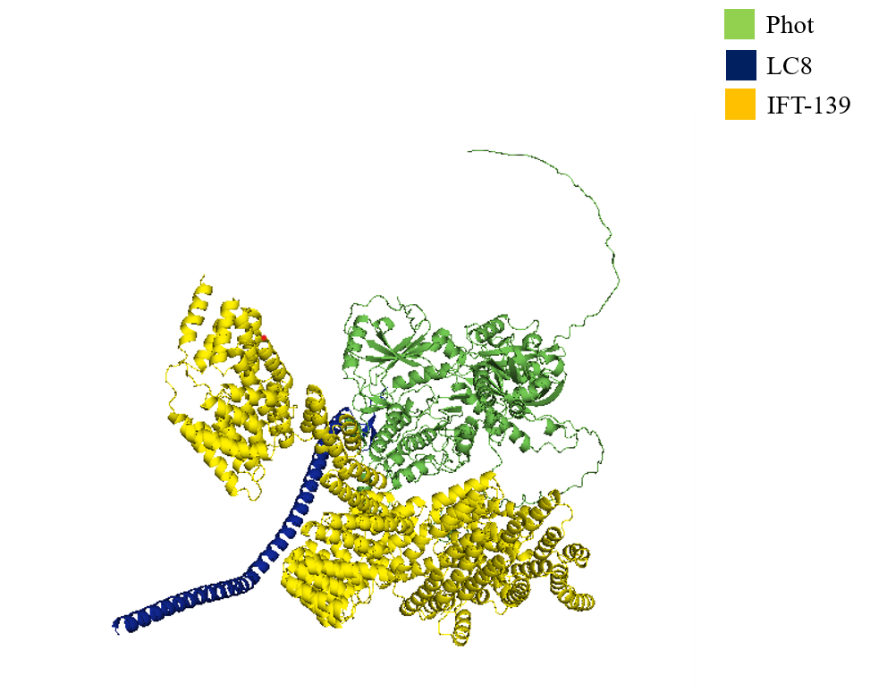

Fig. S1e: Predicted structure showing close vicinity between phototropin, and IFT A subcomplex components IFT 139 and LC-8 of *Klebsormidium nitens*. Structures were predicted using Alpha fold server <https://alphafoldserver.com/> and visualised using Pymol software.

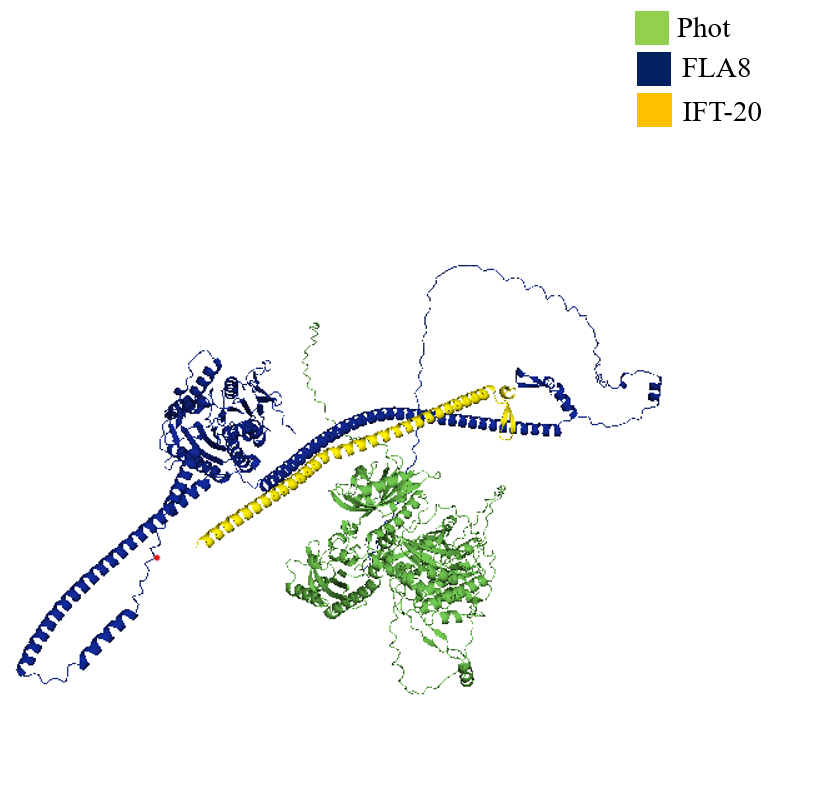

Fig. S1f: Predicted structure showing close vicinity between phototropin, and IFT B subcomplex components IFT20 and FLA8 of *Klebsormidium nitens*. Structures were predicted using Alpha fold server <https://alphafoldserver.com/> and visualised using Pymol software.

**Figure S2**

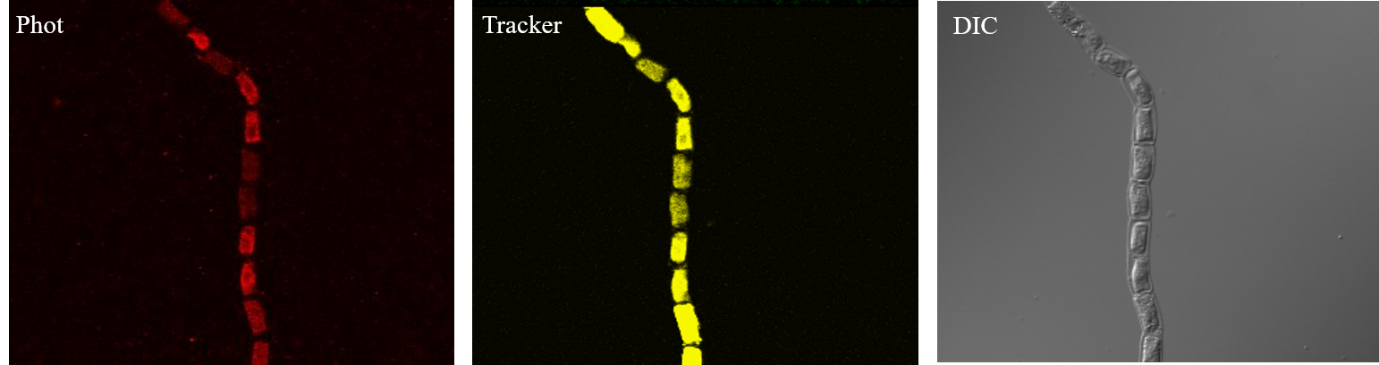

**Fig. S2: Phototropin immunofluorescence multi-cell image along with plasma membrane tracker.**

Immunolocalization of phototropin red channel using primary antibody against Knphot (dilution 1:1000) and secondary antibody anti-rabbit Alexa 546 (1:1000). yellow channel represents signal from plasma membrane tracker.

**Figure S3**

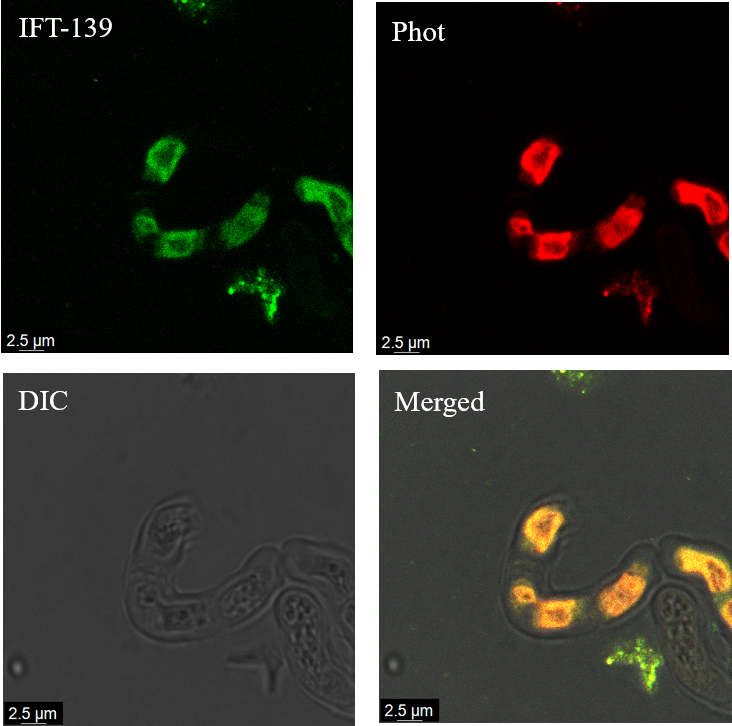

**Fig. S3: Detection and cellular localization of the IFT-A complex component IFT-139 and its colocalization with phototropin in *Klebsormidium nitens*.**

Upper panel in green represents signal of IFT-139 (primary antibody anti-CrIFT139, 1:250), channel in red represents signal for phototropin (primary antibody anti-KnPhot, 1:1000). Lower panel represents DIC and merged image of green channel, red channel and DIC images. Alexa 488 anti-goat and 546 anti-rabbit were used as secondary antibody (1:1000), respectively for green and red channel.

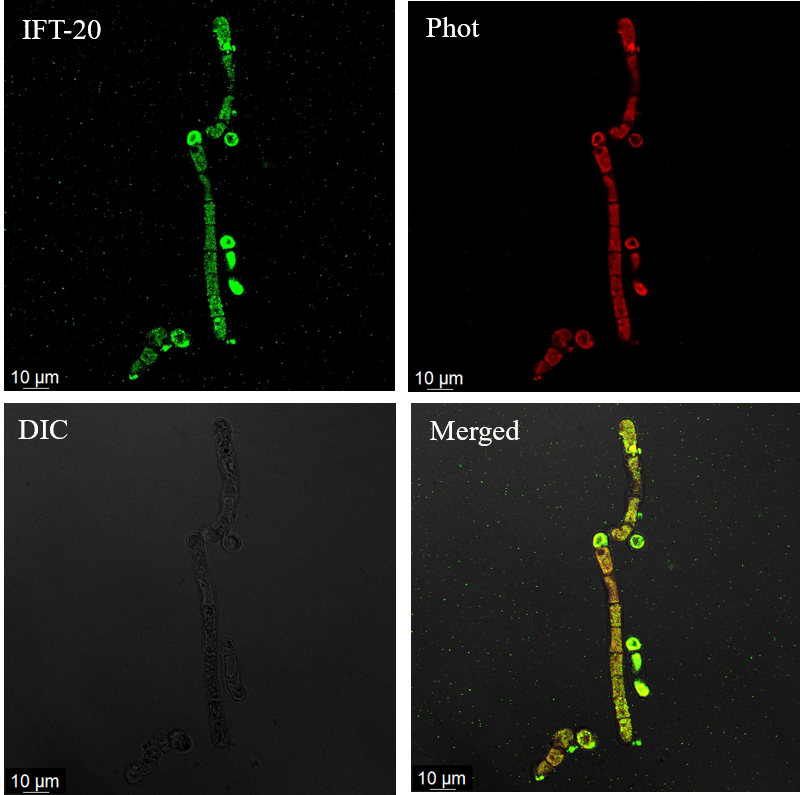
**Figure S4**

**Fig. S4: Detection and cellular localization of the IFT-B complex component IFT-20 and its colocalization with phototropin in *Klebsormidium nitens*.**

Upper panel in green represents signal of IFT-20 (primary antibody anti-CrIFT-20, 1:250), channel in red represents signal for phototropin (primary antibody anti-KnPhot, 1:1000). Lower panel represents DIC and merged image of green channel, red channel and DIC images. Alexa 488 anti-goat and 546 anti-rabbit were used as secondary antibody (1:1000), respectively for green and red channel.

**Figure S5**

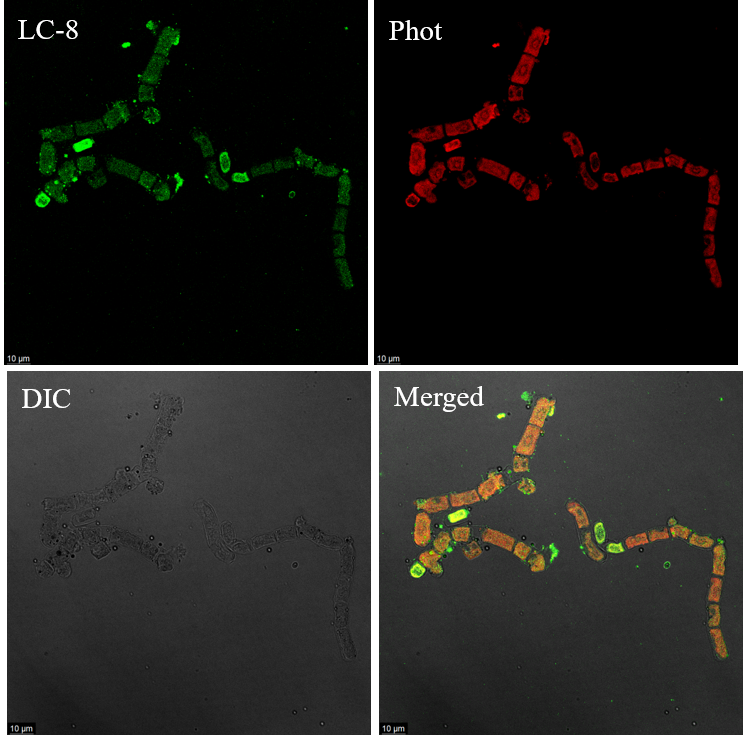

**Fig. S5: Detection and cellular localization of the anterograde motor protein LC-8 and its colocalization with phototropin in *Klebsormidium nitens.***

Upper panel in green represents signal of LC8 (primary antibody anti-CrLC8, 1:250), channel in red represents signal for phototropin (primary antibody anti-KnPhot, 1:1000). Lower panel represents DIC and merged image of green channel, red channel and DIC images. Alexa 488 anti-goat and 546 anti-rabbit were used as secondary antibody (1:1000), respectively for green and red channel.

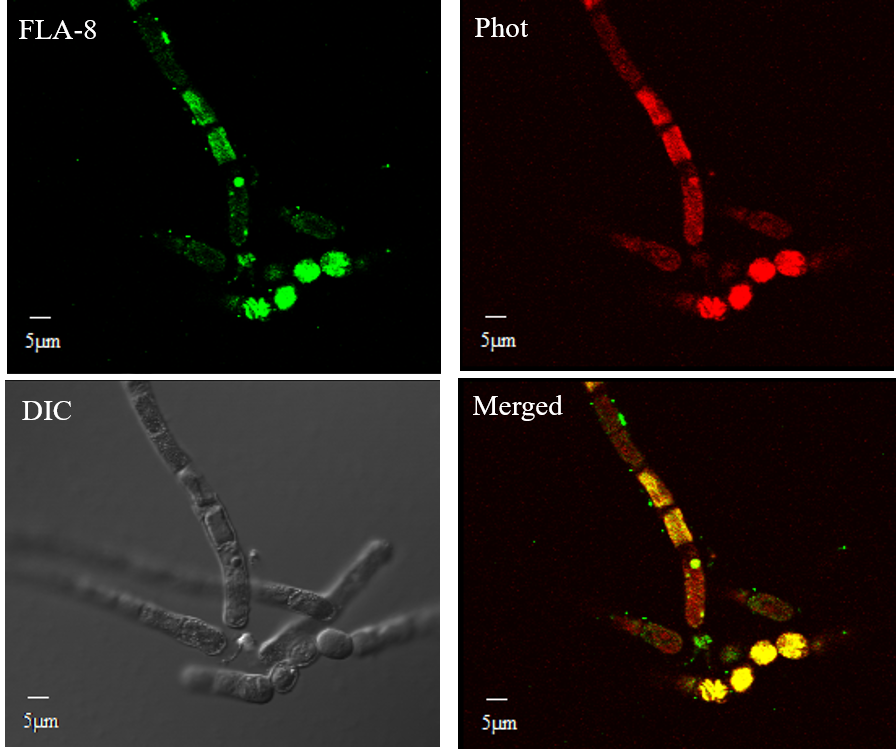
**Figure S6**

**Fig S6: Detection and cellular localization of the retrograde motor protein FLA-8 and its colocalization with phototropin in *Klebsormidium nitens*.**

Upper panel in green represents signal of FLA8 (primary antibody anti-CrFLA8, 1:250), channel in red represents signal for phototropin (primary antibody anti-KnPhot, 1:1000). Lower panel represents DIC and merged image of green channel, red channel and DIC images. Alexa 488 anti-goat and 546 anti-rabbit were used as secondary antibody (1:1000), respectively for green and red channel.

**Figure S7**

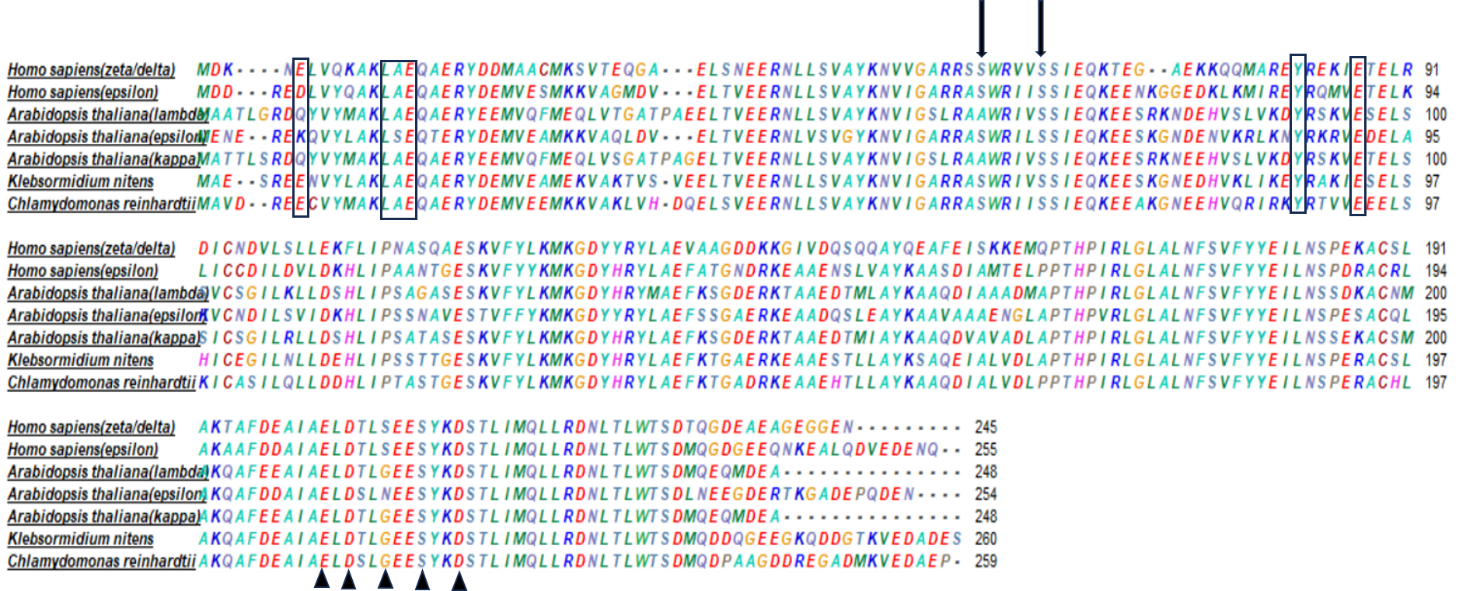

**Fig. S7: Sequence alignment of regulatory protein, *Klebsormidium nitens* 14-3-3 with higher organisms and *Chlamydomonas reinhardtii* algae** using clustal omega and visualisation in Bioedit. Amino acids in box shows dimerization sites, black arrows denote putative phosphorylation site and triangles show EF hand calcium binding motifs.

**Figure S8**

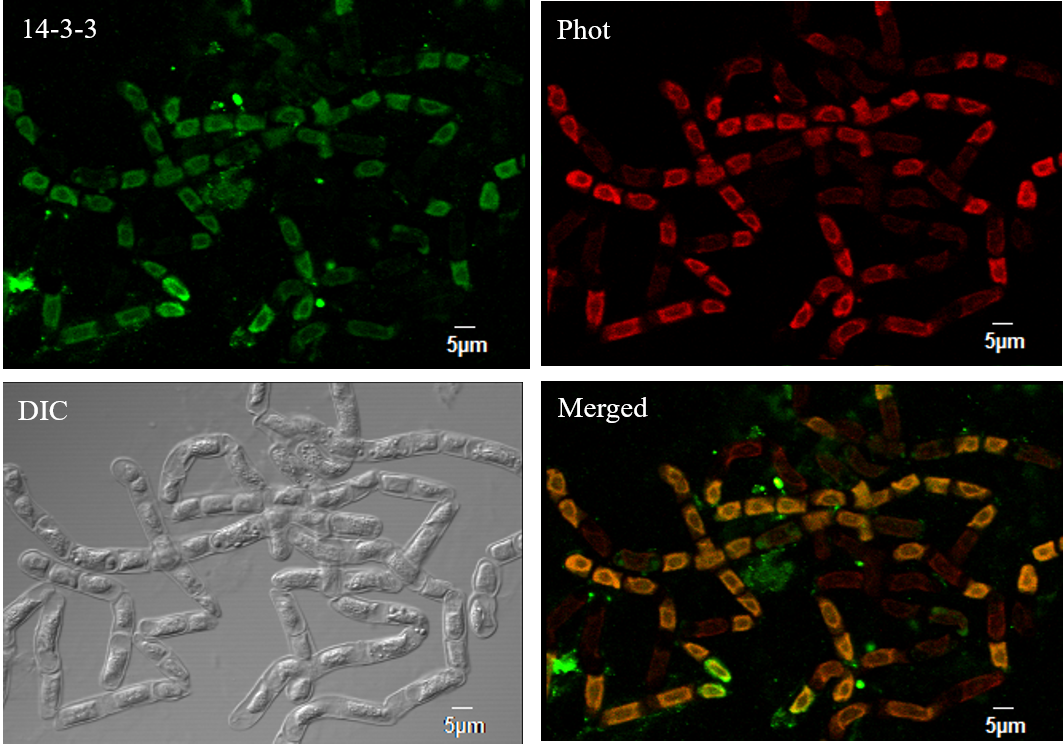

**Fig. S8: Cellular localisation of 14-3-3 and its co-localisation with phototropin in *Klebsormidium nitens***

Upper panel in green represents signal of 14-3-3 (primary antibody anti-Cr14-3-3, 1:250), channel in red represents signal for phototropin (primary antibody anti-KnPhot, 1:1000). Lower panel represents DIC and merged image of green channel, red channel and DIC images. Alexa 488 anti-goat and 546 anti-rabbit were used as secondary antibody (1:1000), respectively for green and red channel.
